# Structural insights into viral genome replication by the severe fever with thrombocytopenia syndrome virus L protein

**DOI:** 10.1101/2022.08.25.505333

**Authors:** Harry M. Williams, Sigurdur R. Thorkelsson, Dominik Vogel, Morlin Milewski, Carola Busch, Stephen Cusack, Kay Grünewald, Emmanuelle R.J. Quemin, Maria Rosenthal

## Abstract

Severe fever with thrombocytopenia syndrome virus (SFTSV) is a phenuivirus that has rapidly become endemic in several East Asian countries. The large (L) protein of SFTSV, which includes the RNA-dependent RNA polymerase (RdRp), is responsible for catalysing viral genome replication and transcription. Here, we present 5 cryo-electron microscopy (cryo-EM) structures of the L protein in several states of the genome replication process, from pre-initiation to late-stage elongation, at a resolution of up to 2.6 Å. We identify how the L protein binds the 5′ viral RNA in a hook-like conformation and show how the distal 5′ and 3′ RNA ends form a duplex positioning the 3′ RNA terminus in the RdRp active site ready for initiation. We also observe the L protein stalled in the early- and late-stages of elongation with the RdRp core accommodating a 9-bp product-template duplex. This duplex ultimately splits with the template binding to a designated 3′ secondary binding site. The structural data and observations are complemented by *in vitro* biochemical and cell-based mini-replicon assays. Altogether, our data provide novel key insights into the mechanism of viral genome replication by the SFTSV L protein and will aid drug development against segmented negative-strand RNA viruses.

## INTRODUCTION

Severe fever with thrombocytopenia syndrome virus (SFTSV) is a segmented negative-strand RNA virus (sNSV) that belongs to the *Phenuiviridae* family in the order *Bunyavirales* [1]. SFTSV was first isolated in China in 2009 [2]. Since then, SFTSV has been identified in several other East Asian countries, including Japan [3], South Korea [4], Vietnam [5], and Taiwan [6]. In addition to humans, SFTSV is known to infect a range of other mammalian hosts, including domesticated animals, such as cats and dogs, as well as wild animals like rodents, boar, and deer [7–9]. SFTSV is thought to be predominantly transmitted via tick bite. However, there is some evidence of human-to-human transmission [2,10,11]. For these reasons, SFTSV is considered a significant threat to global public health but antivirals or other therapeutic approaches and countermeasures are lacking.

A key target for the development of antiviral strategies is the bunyaviral large (L) protein, which includes the viral RNA-dependent RNA-polymerase (RdRp) [12]. The SFTSV L protein is approximately 240 kDa in size and responsible for catalysing viral genome replication and transcription. SFTSV particles contain three genome segments of singe-stranded RNA with negative polarity, denoted S, M and L-segment. During genome replication, negative-sense viral RNA (vRNA) is copied initially into positive-sense complementary RNA (cRNA), which is then used as a template to produce progeny vRNA [12]. Several bunyaviruses are known to initiate genome replication *de novo* by internally producing a short primer, which is subsequently realigned to the 3′ template extremity via a prime-and-realign mechanism. For example, in the case of La Crosse virus (LACV), belonging to the *Peribunyaviridae* family, cRNA synthesis is initiated internally at position 4 of the 3′ vRNA end and the produced short primer is then realigned to the 3′ terminus [13, 14]. In contrast, for Lassa virus (LASV), which is a member of the *Arenaviridae* family, cRNA synthesis is also initiated internally but primer realignment results in an extra G nucleotide at the 5′ end of the vRNA and cRNA [15]. An alternative mechanism involves initiating genome replication de novo terminally without the need for realignment in a similar fashion to that seen for influenza virus [16]. For SFTSV, biochemical data explain terminal initiation and internal initiation equally well and so further studies are needed before concluding whether the SFTSV L protein uses either or both modes to initiate genome replication [17]. Bunyavirus transcription is initiated via a process known as cap-snatching, where the bunyavirus L protein binds to and cleaves off capped fragments of cellular mRNA and uses these as capped primers. This process relies on the endonuclease and a cap-binding activity of the L protein in addition to the RdRp core [18].

Individual endonuclease and cap-binding domains of different bunyaviral L proteins have been investigated by X-ray crystallography [12,17,19–27]. Recently, cryo-electron microscopy (cryo-EM) data from single-particle analysis became available describing relevant functional states of Machupo virus (MACV), LACV and LASV L proteins in complex with vRNA, emphasizing how these complex molecular machines undergo major conformational changes during viral genome replication and transcription [13,14,28,29]. However, to date, structures of the SFTSV and Rift valley fever virus (RVFV) L proteins, as models for phenuiviruses, are only available in an apo state, i.e., without viral RNA [17,30–34]. These structures demonstrated that the overall architecture of the phenuivirus L protein was similar to the structure of the heterotrimeric influenza virus polymerase complex and the peribunyavirus LACV L protein but also identified some notable differences between bunyavirus and influenza virus polymerases, the most apparent being differences in the fingertip’s insertion (SFTSV L protein residues 913 – 920). The fingertip’s insertion is mostly conserved among bunyaviruses [35], presumably because it contains part of the RdRp active site motif F. However, it curiously adopts an alternative conformation in SFTSV compared to LACV L. While representing important first steps, these apo structures of the SFTSV and RVFV L proteins have not yet allowed for an in-depth analysis of protein-RNA interactions: like the structure of the 5′ hook or the mechanism of prime and realign [17]. Such information is however crucial for our understanding of the structure-function relationships underlying viral genome replication and transcription, which is important for antiviral drug development against emerging bunyaviruses.

To address this gap, we took a stepwise approach using a combination of biochemical *in vitro* assays and cryo-EM to visualise and further understand how the structure of the SFTSV L protein changes upon RNA binding and how the L protein catalyses viral genome replication. Thus, here we present 5 cryo-EM structures of the SFTSV L protein in different biologically relevant states from pre-initiation to late-stage of elongation, which are refined to resolutions between 2.6 and 3.5 Å. This structural data is validated by testing selected mutants in a cell-based mini-replicon system. Ultimately, this data provides detailed insights into the mechanism of genome replication by the SFTSV L protein as well as the basis for comparison with other negative-strand RNA viruses like Influenza virus and other bunyaviruses.

## MATERIAL AND METHODS

### Expression and purification of SFTSV L protein

The L gene of SFTSV AH 12 (accession no. HQ116417), which contains a C-terminal StrepII-tag, was chemically synthesised (Centic Biotech, Germany) and cloned into an altered pFastBacHT B vector [17]. A point mutation (D112A) was introduced by mutagenic PCR to reduce the SFTSV L proteins intrinsic endonuclease activity [24]. DH10EMBacY E. coli cells were used to generate recombinant baculoviruses, which were subsequently used for protein expression in Hi5 insect cells. The harvested Hi5 insect cells were resuspended in Buffer A (50 mM HEPES[NaOH] pH 7.0, 1 M NaCl, 10% [w/v] Glycerol and 2 mM DTT), supplemented with 0.05% (v/v) Tween20 and protease inhibitors (Roche, cOmplete mini), lysed by sonication, and centrifuged twice (20,000 x g for 30 minutes at 4 °C). Soluble protein was loaded on Strep-TactinXT beads (IBA Lifesciences) and eluted with 50 mM Biotin (Applichem) in Buffer B (50 mM HEPES[NaOH] pH 7.0, 500 mM NaCl, 10% [w/v] Glycerol and 2 mM DTT). L protein-containing fractions were pooled and diluted 1:1 with Buffer C (20 mM HEPES[NaOH] pH 7.0) before loading on a heparin column (HiTrap Heparin HP, Cytiva). Proteins were eluted with Buffer A and concentrated using centrifugal filter units (Amicon Ultra, 30 kDa MWCO). The proteins were further purified by size-exclusion chromatography using a Superdex 200 column (Cytiva, formerly GE Life Sciences) in buffer B. Purified L proteins were concentrated as described above, flash frozen and stored at −80 °C.

### Polymerase assay

For the standard polymerase assay, 0.9 μM of the SFTSV L protein was incubated sequentially with 1.8 μM of single-stranded 5′ promoter RNA and 1.8 μM of single-stranded 3′ promoter RNA in assay buffer (100 mM HEPES, 50 mM KCl, 2.5 mM MgCl_2_, and 2 mM DTT, pH 7) on ice for 20 minutes (Supplementary Table 1). For primer-dependent reactions, 18 μM of the respective primer was added (making for a 10-fold molar excess to promoter RNA) to SFTSV L protein bound to promoter RNA and the mix incubated for an additional 20 minutes on ice. The polymerase reaction was started by the addition of NTPs (0.2 mM ATP/CTP/UTP and 0.1 mM GTP spiked with 166 nM, 5 µCi[α]^32^P-GTP, unless stated otherwise) in a final reaction volume of 10 μL. After incubation at 30 °C for 2 hours, the reaction was stopped by adding an equivalent volume of RNA loading buffer (98% formamide, 18 mM EDTA, 0.025 mM SDS, xylene cyanol and bromophenol blue) and heating the samples at 95 °C for 5 minutes. Products were then separated by denaturing gel electrophoresis using 25% polyacrylamide 7 M urea gels and 0.5-fold Tris-borate-EDTA running buffer. Signals were visualised by phosphor screen autoradiography using a Typhoon FLA-7000 phosphorimager (Fujifilm) operated with the FLA-7000 software.

### Rift valley fever virus mini-replicon system

The experiments were performed using the RVFV mini-replicon system described previously [36]. L genes were amplified using mutagenic PCR from a pCITE2a-L template to produce either wild-type or mutated L gene expression cassettes. PCR products were gel purified and quantified spectrophotometrically. All mutations were confirmed by sequencing of the PCR products. BSR-T7 cells were transfected per well of a 24 well plate with 250 ng of L gene PCR product, 500 ng of pCITE expressing NP, 750 ng of pCITE expressing the mini-genome RNA encoding Renilla luciferase (Ren-Luc), and 10 ng of pCITE expressing the firefly luciferase as an internal control, all under control of a T7 promoter. For transfection we used Lipofectamine 2000 according to the manufacturer’s instructions. At 24 hours post-transfection, either total cellular RNA was extracted for Northern blot analysis using an RNeasy Mini kit (Qiagen) or cells were lysed in 100 μL of passive lysis buffer (Promega) per well, and firefly luciferase and Ren-Luc activity quantified using the dual-luciferase reporter assay system (Promega). Ren-Luc levels were corrected with the firefly luciferase levels (resulting in standardised relative light units [sRLU]) to compensate for differences in transfection efficiency or cell density. Data were evaluated in Prism and are always presented as the mean of a given amount (n) of biological replicates as well as the respective standard deviation (SD). To confirm general expressibility of the L protein mutants in BSR-T7 cells, the cells were transfected with 500 ng L gene PCR product expressing C-terminally 3xFLAG-tagged L protein mutants per well. Cells were additionally infected with Modified Vaccinia virus Ankara expressing a T7 RNA polymerase (MVA-T7) to boost the expression levels and, thereby facilitate detection by immunoblotting. At 24 hours post-transfection, cells were lysed in 50 μL passive lysis buffer (Promega) per well. After cell lysis and separation in a 3-8% Tris-acetate polyacrylamide gel (Invitrogen), proteins were transferred to a nitrocellulose membrane (Cytiva). FLAG-tagged L protein mutants were detected using horseradish peroxidase-conjugated anti-FLAG M2 antibody (1:9,000) (A8592; Sigma-Aldrich) and bands were visualised by chemiluminescence using Clarity Max Western ECL Substrate (Bio-Rad) and a FUSION SL image acquisition system (Vilber Lourmat).

### Northern blot analysis

For the Northern blot analysis, 600 ng of total cellular RNA was separated in a 1.5% agarose-formaldehyde gel and transferred onto a Roti-Nylon plus membrane (pore size 0.45 μΜ, Carl Roth). After UV cross-linking and methylene blue staining to visualise 28 S rRNA, the blots were hybridised with a 32P-labelled riboprobe targeting the Ren-Luc gene. Transcripts of the Ren-Luc gene and complementary replication intermediate RNA of the minigenome were visualised by autoradiography using a Typhoon FLA-7000 phosphorimager (Fujifilm) operated with the FLA-7000 software. Quantification of signals for antigenomic RNA and mRNA was done in Fiji.

### Sample preparation for cryo-EM

#### 5′ HOOK structure

The SFTSV L protein at a concentration of 3 μM in assay buffer (100 mM HEPES, 50 mM KCl, 2.5 mM MgCl_2_, and 2 mM DTT, pH 7) was mixed with single-stranded L 5′ (1-20) cRNA in a 2-fold molar excess. The mixture was incubated on ice for 30 minutes and then centrifuged at 15,000 x g for 10 minutes at 4 °C. Aliquots of 3 μL were applied to Quantifoil R 2/1 Au G200F4 grids, immediately blotted for 2 seconds, and plunge frozen into liquid ethane/propane cooled to liquid nitrogen temperature using a FEI Vitrobot Mark IV (4 °C, 100% humidity, blotting force –10).

### EARLY-ELONGATION, EARLY-ELONGATION-ENDO, and LATE-ELONGATION structures

The SFTSV L protein at a concentration of 3 μM in assay buffer (100 mM HEPES, 50 mM KCl, 2.5 mM MgCl_2_, and 2 mM DTT, pH 7) was mixed sequentially with single-stranded L 5′ (1-20) cRNA and single-stranded L 3′ (1-20) cRNA modified with 6 additional A bases, in a 2-fold molar excess. After 30 minutes of incubation on ice, the reaction was initiated by addition of NTPs (0.63 mM ATP/CTP/GTP/nhUTP). After incubation at 30 °C for 2 hours, the sample was centrifuged at 15,000 x g for 10 minutes at 4 °C. Aliquots of 3 μL were applied to Quantifoil R 2/1 Au G200F4 grids, immediately blotted for 2 seconds, and plunge frozen into liquid ethane/propane cooled to liquid nitrogen temperature using a FEI Vitrobot Mark IV (4 °C, 100% humidity, blotting force –10).

### RESTING structure

The SFTSV L protein at a concentration of 3 μM in assay buffer (100 mM HEPES, 50 mM KCl, 2.5 mM MgCl_2_, and 2 mM DTT, pH 7) was mixed with single-stranded L 5′ (1-20) cRNA and single-stranded L 3′ (1-20) cRNA in a 2-fold molar excess. After 30 minutes of incubation on ice, the reaction was supplemented with a short 2 nt primer (AC), which was at a 1:1 molar ratio with the RNA species. After incubation at 30 °C for 2 hours, the sample was then centrifuged at 15,000 x g for 10 minutes at 4 °C. Aliquots of 3 μL were applied to Quantifoil R 2/1 Au G200F4 grids, immediately blotted for 2 seconds, and plunge frozen into liquid ethane/propane cooled to liquid nitrogen temperature using a FEI Vitrobot Mark IV (4 °C, 100% humidity, blotting force –10).

### Cryo-EM single-particle analysis

#### Data collection

Grids were loaded into a 300-kV Titan Krios transmission electron microscope (Thermo Scientific) equipped with a K3 direct electron detector and a post-column GIF BioQuantum energy filter (Gatan). Micrograph movies were typically collected using the EPU software (Thermo Scientific) at a nominal magnification of 105,000x with a pixel size of 0.85 Å using a defocus range of −0.8 to −2.6 μm, unless stated otherwise. The sample was exposed with a dose of 15 electrons per physical pixel per second (estimated over vacuum) for 2.5 s and thus, a total dose of 50 electrons per Å^2^. Recorded movie frames were fractionated into 50 frames. A summary of the key data collection statistics for each cryo-EM structure is provided in Supplementary Table 2.

#### Image processing

All movie frames were imported in RELION 3.1 [37] or RELION 4.0 [38] and motion corrected using its own implementation of MotionCor2 [39]. Particles were picked automatically using the pre-trained BoxNet in Warp [40] and the calculation of contrast transfer function (CTF) estimation and correction was done in RELION with CTFFIND4 [41]. Picked particles were extracted and processed in RELION 3.1/4.0 with visual inspection done with UCSF Chimera [42]. First, particles were subjected to 2D classification at 4x binned pixel size typically as an initial check of the quality of the dataset. Then, all particles were used for 3D classification in an iterative manner using a 50 Å low pass filtered map from apo SFTSV L protein [17]. Incomplete, low resolution or 3D classes containing damaged particles were excluded from further data analyses. The remaining particles were re-extracted with the original pixel size value for further 3D classification and refinement as described for the 5′ HOOK structure (Supplementary Figure 1 – 2); EARLY-ELONGATION, EARLY-ELONGATION-ENDO, and LATE-ELONGATION structures (Supplementary Figure 3 – 4); and, RESTING structure (Supplementary Figure 5 – 6). All cryo-EM maps were CTF refined and Bayesian polished before final refinement in RELION except the RESTING structure which, after 3D classification, was imported as a particle stack in cryoSPARC [43] where final refinement was done with non-uniform refinement [44].

All final cryo-EM density maps were generated by the post-processing feature in RELION and sharpened or blurred into MTZ format using CCP-EM [45] except the EARLY-ELONGATION-ENDO structure where the final map was sharpened using DeepEMhancer [46] and the RESTING structure which was sharpened using the local sharpening function of cryoSPARC. The resolutions of the cryo-EM density maps were estimated at the 0.143 gold standard Fourier Shell Correlation (FSC) cut-off. A local resolution was calculated using RELION 3.1/4.0 except the RESTING structure which was calculated in cryoSPARC and displayed with Chimera visualisation tools. A summary of the key processing statistics for each cryo-EM structure is provided in Supplementary Table 2.

#### Model building, refinement, and validation

The published corrected apo structure of the SFTSV L protein [30] was used as the starting point for the RNA-bound structures published in this study which then underwent iterative rounds of model building in Coot [47] and real-space refinement in Phenix [48]. The comprehensive validation tool in Phenix was used to assess model quality [49, 50]. Coordinates and EM maps are deposited in the EMDB and PDB databases as indicated in Table 1. Figures have been generated using CCP4mg [51], PyMOL (Schrödinger), and UCSF ChimeraX [52]. A summary of the key refinement and validation statistics for each cryo-EM structure is provided in Supplementary Table 2.

**Table 1.** Overview of Cryo-EM Structures. An overview of the cryo-EM structures is provided with the reported resolution (Å), any RNA ligands present, PDB and EMDB accession codes, as well as supporting comments highlighting the key features of each structure.

| Name | Res (Å) | RNA Ligands | PDB Code | EMDB Code | Comment |
| --- | --- | --- | --- | --- | --- |
| 5' HOOK | 3.4 | L 5' (1-20) | 8AS6 | EMD-15607 | Only 5' cRNA is bound to the L protein in the hook-like motif. The L protein core is clear and motif F is unstructured. |
| EARLY-ELONGATION | 2.6 | L 5' (1-20)<br>L 3' + 6A (1-26) | 8AS7 | EMD-15608 | The 5' and 3' cRNA form a distal duplex structure with the 3' cRNA shown feeding into the L protein core. Short 3 nt product in L protein core. Stalling nhUTP clearly bound at +1 position. |
| EARLY-ELONGATION-ENDO | 3.6 | L 5' (1-20)<br>L 3' + 6A (1-26) | 8ASB | EMD-15610 | The 5' and 3' cRNA form a distal duplex structure with the 3' cRNA shown feeding into the L protein core. There is also a short 3 nt-long product in L protein core. However, in this structure, we see the endonuclease in a raised conformation unlike that seen elsewhere. Stalling nhUTP clearly bound at +1 position. |
| LATE-ELONGATION | 2.6 | L 5' (1-20)<br>L 3' + 6A (1-26) | 8ASD | EMD-15614 | The distal duplex structure is disrupted and there is a long 9-bp product-template duplex in the L protein core. Stalling nhUTP clearly bound at +1 position. The CBD can also be fit with the putative cap-mRNA binding site occupied by the Arginine blocking motif. |
| RESTING | 3.2 | L 5' (1-20)<br>L 3' (1-20)<br>Primer AC (1-2) | 8ASG | EMD-15615 | The 5' cRNA is bound as in the 5' HOOK structure. Excess of 3' cRNA is bound to the 3' secondary binding site. A small number of 3' cRNA nts can be fit in the L protein core but the region of map for the AC primer is not resolved. |

## RESULTS

### Overview

We have determined 5 structures of the SFTSV L protein in several biologically relevant states ranging from pre-initiation to late-stage elongation. An overview of all structures including their key features is provided in Table 1. First, we incubated the L protein with nt 1-20 of the L-segment 5′ cRNA (5′ HOOK, 3.4 Å) (Supplementary Table 1; Supplementary Figures 1 and 2). This initial structure enabled us to identify the pocket in which the L protein coordinates the 5′ RNA terminus in a so-called hook-like conformation, which is known to be important for activation of the polymerase function [17].

To stall the L protein in an active genome replication mode, we incubated the L protein with nts 1-20 of the L-segment 5′ cRNA and a modified 3′ cRNA template with 6 additional A’s (Supplementary Table 1). These were incubated together in the presence of ATP, CTP, and GTP spiked with non-hydrolysable UTP (nhUTP) (Supplementary Figures 3 and 4). With this, we observed three elongation-state structures: two in which the L protein is stalled at position G4 of the modified 3′ cRNA template (EARLY-ELONGATION, 2.6 Å and EARLY-ELONGATION-ENDO, 3.6 Å) and another in which the L protein is stalled at position A21 of the modified 3′ cRNA template (LATE-ELONGATION, 2.6 Å). While the binding of viral RNAs in the two early-elongation structures appear similar with the distal 5′ cRNA region forming a duplex with the 3′ cRNA template, there is a key difference in the overall conformation of the L protein. In the EARLY-ELONGATION-ENDO structure, the endonuclease domain adopts an entirely different ‘raised’ conformation by comparison to the endonuclease domain in the EARLY-ELONGATION structure, which is broadly similar to that seen in the apo and LATE-ELONGATION structures. In the latter, the CBD can also be fitted and the putative cap-binding site appears blocked by the arginine blocking motif previously described [30–33]. Therefore, in total, we obtained 3 structures of the L protein stalled in active genome replication mode: two stalled in the early-stages of elongation, and a final structure stalled in a late-stage of elongation.

To identify the structural basis for replication priming, we incubated the L protein with nts 1-20 of the L-segment 5′ cRNA and 3′ cRNA termini and added a short 2 nt (AC) primer (Supplementary Table 1; Supplementary Figures 5 and 6). Processing of data collected from this grid led to a single structure (RESTING, 3.2 Å), in which the 5′ RNA hook binds in the same way as that seen in our other structures. However, rather than formation of the distal duplex and the 3′ cRNA template entering the active site, we observe the 3′ cRNA terminus binding to an alternative region, denoted the 3′ secondary binding site in analogy to similar binding sites identified in LASV [28] and LACV L proteins [13] and also the influenza virus polymerase complex [53]. The distal duplex is not formed in this structure and we observe a weak signal in this region of the map that likely corresponds to an average of the RNA strands in multiple different conformations. Nevertheless, this structure demonstrates that both the 3′ and 5′ RNA termini can be bound simultaneously to the hook-binding and secondary binding sites, respectively. We propose that this might represent a resting conformation after finishing one round of genome replication and just before another pre-initiation complex is formed with the distal duplex stabilizing the 3′ RNA and guiding it into the RdRp active site for initiation. However, in this structure, we can also confidently fit 2 nts of the 3′ cRNA in the active site, which indicates that the observed map represents a mixture of resting and pre-initiation states, which would explain the relative lack of definition in some regions of the map, making it challenging to interpret.

### The L protein during pre-initiation of viral genome replication

The 5′ HOOK structure demonstrates that the SFTSV L protein undergoes several conformational changes as a result of binding to the terminal 20 nts of the 5′ cRNA in comparison to the apo structure (PDB: 6L42, 6Y6K, and 7ALP) [30–33]. The most apparent changes being an opening up of the vRNA binding lobe (vRBL), the neighbouring fingers domain, and the PA-C like core lobe, allowing the 5′ RNA to bind in a positively charged cleft (Figure 1A). This cleft is covered by a short 3-turn α-helix (M437 – H447) and supported on the periphery by a short loop (H1038 – D1046) (Figure 1A). Of note, both of these structural motifs were missing in the published apo structures, suggesting here that the 5′ RNA has a role in stabilising this region of the L protein.

**Figure 1.**
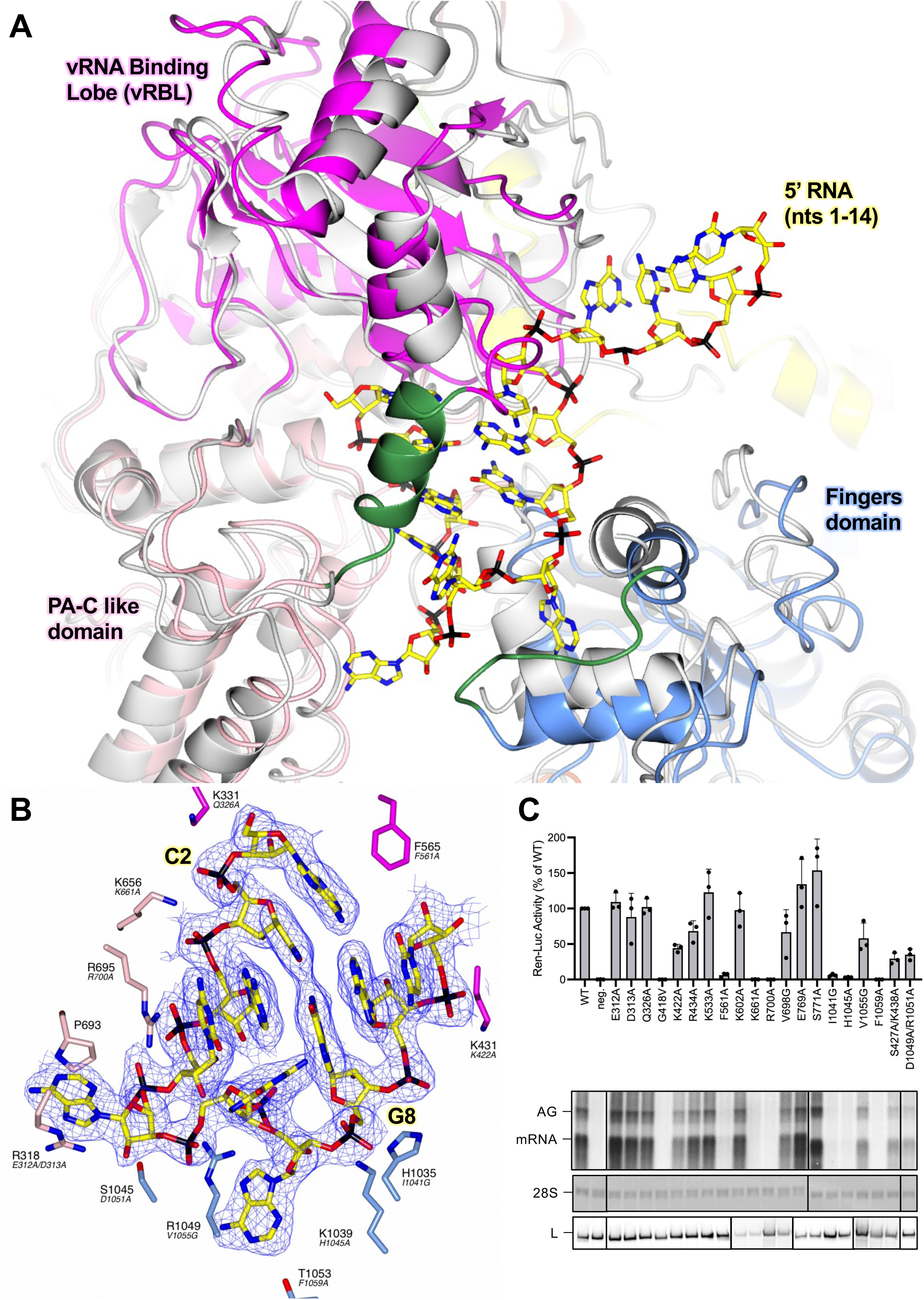
Conformational changes following the binding of the 5′ cRNA hook. (A) The SFTSV L protein (5′ HOOK) is shown with the published apo structure of the SFTSV L protein (PDB: 7ALP, backbone coloured grey) superposed by SSM in CCP4mg [51]. A short 3-turn α-helix (M437 – H447) and loop (H1038 – D1046), which are both stabilised by the binding of the 5′ cRNA nts, are shown in green. (B) A close-up of the SFTSV L protein hook-binding site is shown with the EARLY-ELONGATION map overlaid. Bound 5′ cRNA is shown and coloured yellow with the exception of C2 and G8, which together form the only cognate base-base interaction and are labelled. Residues from the SFTSV L protein are labelled and coloured according to the assigned domain (pink for the PA-C like-domain, blue for the fingers, and magenta for the vRBL) with the corresponding RVFV L protein residue shown in black italicised text underneath. Not all hook-interacting residues are shown for clarity. (C) RVFV mini-replicon data for L protein with mutations in the proposed 5′ hook binding site presenting luciferase reporter activity (in standardised relative light units relative to the wild-type L protein (WT)). Data were presented as mean values ± SD of three biological replicates (n = 3). All biological replicates are shown as black dots (top panel). Middle panels present Northern blotting results with signals for antigenomic viral RNA (AG, equal to cRNA), viral mRNA (mRNA) and 28 S ribosomal RNA (28 S) as a loading control, and the bottom panel shows Western blot detection of FLAG-tagged L proteins (L) to demonstrate general expressability of the mutants. Uncropped original blots/gels are provided in the Supplementary Data.

### Structure of the 5′ cRNA hook

We find the 5′ cRNA bound in each of our L protein structures and, depending on the structure, we can fit 11 – 17 nts into the corresponding map region. From this data, we found that the first 10 nts of the SFTSV 5′ RNA form the characteristic hook-like motif (Figure 1B). In addition, in our structures, the 5′ RNA hook only appears to be stabilised by one canonical base-base interaction between C2 and G8. The SFTSV hook structure hence appears to be closer to that of LASV, which is also stabilised by one canonical base-base interaction [28], in comparison to that of LACV, which is stabilised by two canonical base-base interactions [14] (Supplementary Figure 7). In addition to the C2/G8 base-pairing interaction, amino acid sidechains from the fingers, finger node, and the core lobe contribute towards 5′ RNA hook coordination.

For example, the K656 sidechain is shown to interact with the backbone phosphates of nts C2 and A3 (Figure 1B). The N765 sidechain also interacts with nt A3, with the sidechain amino group coordinating one of the A3 base nitrogens, while the carboxyl group interacts with the A3 ribose O2’ hydroxyl. There are two sidechain interactions stabilising nt C4: R1049 and H447. These coordinate the C4 ribose O2’ hydroxyl and C4 base oxygen, respectively. The R695 sidechain also stacks against the C4 base providing additional stability. The A5 base is sandwiched between the R318 and P693 sidechains. Similarly, there are not many interactions visible stabilising the G6 base, only the H447 sidechain coordinating the G6 ribose oxygen (Supplementary Figure 8A). In addition to this interaction, in our EARLY-ELONGATION structure, we noticed an interesting ring-shaped blob of density between the G6 base and the adjacent R1043 sidechain (Supplementary Figure 8B). After exploring several different options, including individual NTPs bound non-specifically at this position, we concluded that the best fit was a single HEPES molecule, which is the buffer used during the L protein/cRNA incubation step. Here, the HEPES ring is shown sandwiched in between the G6 base and the R1043 sidechain (Supplementary Figure 8B). The base A7 is shown to sit comfortably in a relatively tight pocket delineated by the sidechains of F1032, L1036, F1048, R1049, and M1052 (Supplementary Figure 8C). Here, the R1049 sidechain also coordinates the A7 backbone phosphate. A9 is coordinated by the S562 and H563 sidechains, which interact with the A9 O2’ hydroxyl and a base nitrogen atom, respectively (Supplementary Figure 8D). Finally, the C10 base is sandwiched between the A9 base and the K431 sidechain. There is then a sharp turn between nt 10 and 11 of the 5′ RNA marking the end of the hook motif.

### Mutational analysis of 5′ RNA hook-binding residues

To investigate the importance of the key 5′ RNA hook-binding residues, we mutated 21 different amino acids of the L protein (18 single, 2 double-mutants) and tested for genome replication and transcription activity in a viral mini-replicon system. In this system, genomic viral RNA is produced by the T7 polymerase from a plasmid and then serves as a template for viral genome replication and transcription, producing Ren-Luc mRNA, antigenomic viral RNA and subsequently also further genomic viral RNA copies, all catalyzed by L and N proteins [36]. As we had a RVFV mini-replicon system already established, we mapped the identified SFTSV residues onto the closely-related RVFV L protein (Supplementary Figure 9), and then tested the corresponding positions in this RVFV mini-replicon system [36]. Of the 21 mutants tested, 11 showed wild type-like activity (i.e., Ren-Luc activity, >50%) and 3 mutants showed a slightly reduced activity (i.e., Ren-Luc activity, 8-49%) (Supplementary Table 3, Figure 1C). However, mutations of 7 residues led to a strong decrease in RVFV L protein activity (i.e., Ren-Luc activity, <7%), including: G418V (SFTSV: G427), F561A (SFTSV: F565), K661A (SFTSV: K656), R700A (SFTSV: R695), I1041G (SFTSV: H1035), H1045A (SFTSV: K1039), and F1059A (SFTSV: T1053) (Supplementary Table 3, Figure 1C). Of those residues critical for L protein function in the RVFV mini-replicon system, the corresponding K656, R695, and K1039 directly interact with A3, C4/A5, and G8 of the 5′ cRNA in our structure of the SFTSV L protein. In addition, H1035 appears to line the hook binding cleft, whereas T1053 has a long interaction (>4 Å) with the A7 base of the 5′ cRNA, while G427 and F565 are adjacent to the A1 and C10 nts of the 5′ cRNA (Figure 1B). Overall, this analysis suggests that the 5′ RNA binding site is important for both genome replication and transcription as no selective defect in RNA synthesis could be detected. This matches previous observations for LASV L protein [28].

### The transition from pre-initiation to elongation

To capture the SFTSV L protein in an actively replicating state, we incubated the L protein with nts 1-20 of the SFTSV L 5′ cRNA and a 3′ cRNA template modified with 6 additional A’s at the 5′ end together with ATP, GTP, CTP, and nhUTP. The SFTSV L 3′ cRNA has only one A in the first 20 nts and this is at position 19. If we had stalled the L protein using nhUTP with this un-modified 3′ cRNA then the difference between the stalled and un-stalled products would have been only 1 nt and this would have been difficult to visualise biochemically. Thus, we decided to add 6 additional A’s to the 5′ end of the 3′ cRNA boosting the overall size of the template from 20 nts to 26 nts. If the L protein stalled at the first A in the 3′ cRNA template, this would have produced a stalling product at 19 nts, whereas the un-stalled product would now be 26 nts. This greater difference in product size allowed us to visualise the stalling reaction clearly by biochemical means.

After cryo-EM data collection and processing, we identified a major class in which the L protein appeared to be stalled in the later stages of elongation with a 9-bp product-template duplex in the L protein core (LATE-ELONGATION, 2.6 Å; Supplementary Figure 3 - 4). Upon further inspection, we discovered that the L protein had seemingly stalled at A21 of the 3′ cRNA. Using similar conditions, biochemical analysis showed that incorporation of nhUTP into the NTP mix leads to two major products: one at 19 nts, which was the expected stalling product length given that the first A in the template sequence is at position 19, and a second, 21 nt-long product. As it is known that viral RNA polymerases are relatively low-fidelity enzymes [54], we hypothesized that the second major stalling product at 21 nts was probably the result of the L protein misincorporating CTP at A19 in the template before stalling at A21. To test this hypothesis, we titrated CTP out of the NTP mix spiked with nhUTP and found that as we reduced the concentration of CTP, the intensity of the larger, 21 nt band, decreased in a corresponding manner (Figure 2). At the same time, the intensity of the smaller 19 nt band increased (Figure 2). Thus, we could demonstrate both *in vitro* and structurally that the SFTSV L protein can misincorporate CTP in place of UTP. There is also a minor product at ~20 nts (Figure 2). We think this could either be termination after a second misincorporated CTP or possibly the result of misincorporating nhUTP at this position. The second of these two options seems less likely but that may explain why this product has a much weaker intensity. It is also noteworthy that as we titrate CTP out of the NTP mix, the intensity of two smaller bands in the 10 – 15 nt range increases (Figure 2). We think these represent early termination events where the L protein reaches a poly-G sequence in the 3′ cRNA strand and then stalls due to lack of CTP. This would explain why the intensity of these products increases as CTP is titrated out of the NTP mix.

**Figure 2.**
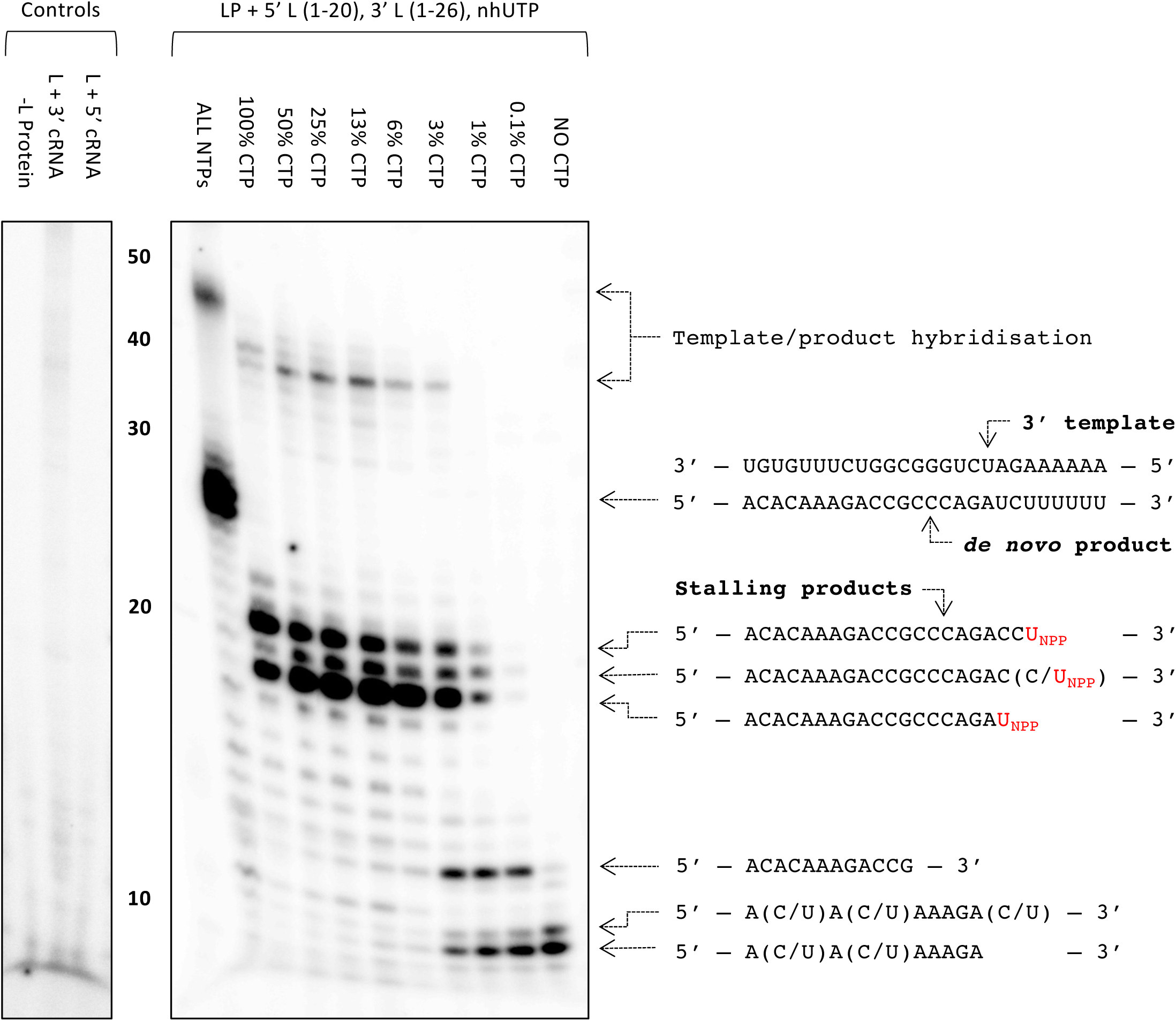
Stalling of the SFTSV L protein and misincorporation of CTP for UTP. The effect of removing CTP from the NTP mix in the presence of nhUTP (U_NPP_) on the polymerase activity of purified SFTSV L protein (LP) was tested *in vitro*. The reactions were carried out with the L 5′ cRNA (nts 1-20) and L 3′ cRNA (nts 1-26) present under standard polymerase assay conditions (see Materials and Methods), with the exception of the NTPs used in the CTP titration conditions, which were diluted 1:10 resulting in a final concentration of 20 µM ATP, UTP and 10 µM GTP spiked with 166 nM, 5 µCi[α]^32^P-GTP. The final concentration of CTP in the NTP mix varied from 20 µM to 0 µM. Products were separated by denaturing gel electrophoresis and visualised by autoradiography. Uncropped original blots/gels are provided in the Supplementary Data.

Processing of the cryo-EM data identified two other major classes with the L protein stalled at a much earlier point in the 3′ cRNA template and with significant differences regarding the endonuclease domain (Supplementary Figures 3 and 4). Whereas in the EARLY-ELONGATION structure the endonuclease adopts a similar conformation to that seen in the published apo structure [30–33], in the EARLY-ELONGATION-ENDO structure, the endonuclease adopts a ‘raised’ conformation (Supplementary Figures 3 and 4). After inspecting the initial maps, we found that in both early elongation structures the 3′ RNA template was positioned with 4 nts into the L protein core (3′-UGUG…-5′), with a short 3 nt product (5′-ACA-3′) and the nhUTP at the +1 position, where we would actually expect CTP. Given that we knew the L protein could misincorporate CTP in place of UTP, we considered it possible the L protein could also misincorporate UTP in place of CTP. If this was true, the L protein could presumably also misincorporate nhUTP leading to the observed early stalling product.

To test this hypothesis, we selected the SFTSV S_9U_ segment vRNA as this genome end has a naturally occurring poly-G sequence. If the L protein can incorporate UTP in place of CTP, then in the absence of CTP, the L protein should be able to read-through the poly-G sequence (nts 11-15) by incorporating UTP. If not, then we should observe a stalling product around 11 nts. We additionally used a 16 nt primer (Primer B-OH, Supplementary Table 1) to increase any stalling product size by 13 nts, accounting for the 3 bp overlap with the template. In our *in vitro* polymerase assay setup, when we remove CTP from the NTP mix but retain UTP, we still see the full-length product (Supplementary Figure 10). When we then also titrate UTP out of the NTP mix, we see the emergence of a product at the approximate size of the expected primed stalling product (Supplementary Figure 10). Together, this strongly suggests that the L protein can also incorporate UTP in place of CTP supporting our hypothesis that the early-stalling structures we observed are likely the result of nhUTP being misincorporated in place of CTP. To conclude, we obtained 3 structures of the SFTSV L protein stalled in actively replicating states: two structures stalled at position G4 of the 3′ cRNA (EARLY-ELONGATION and EARLY-ELONGATION-ENDO), and one structure stalled at position U21 of the 3′ cRNA (LATE-ELONGATION), which we refer to as early- and late-stage elongation states, respectively.

### The L protein early elongation structure

The two early elongation structures, denoted EARLY-ELONGATION and EARLY-ELONGATION-ENDO, demonstrate that when both 5′ and 3′ 1-20 nts cRNAs are present, the SFTSV L protein undergoes further remodelling, mostly involving residues in the thumb and thumb ring domains. These structural changes allow the L protein to accommodate the 5′/3′ distal duplex region, which is shown to curl around the vRBL, adopting a similar conformation to the full promoters observed for LACV [14] and LASV [28] (Figure 3A). However, there is a key difference in the number of canonical base-pair interactions between the 5′ and 3′ RNA strands. Unlike in other related bunyaviruses, the conserved 5′ and 3′ RNA termini of the SFTSV L segment are not particularly complementary with theoretically only 11 matches within the first 20 nts, including only 2 exact matches (nts 13 and 14) in the distal duplex region (nts 11-18) (Supplementary Figure 11). However, in the EARLY-ELONGATION and EARLY-ELONGATION-ENDO structures, we observe 7 base-pair interactions in the distal duplex. This is possible because the 5′ and 3′ RNA strands are shifted by 1 nt in respect to each other, meaning that G11 of the 5′ cRNA base-pairs with C12 of the 3′ cRNA, C12 of the 5′ cRNA base-pairs with G13 of the 3′ cRNA and so on (Figure 3B). In LACV, the 5′/3′ distal duplex is stabilised by 6 base-pair interactions [13], whereas, in LASV there are 8 base-pair interactions [28]. This shift between the 3′ and 5′ RNA strands presumably enables the SFTSV L protein to overcome the general lack of complementarity in the 3′/5′ duplex region.

**Figure 3.**
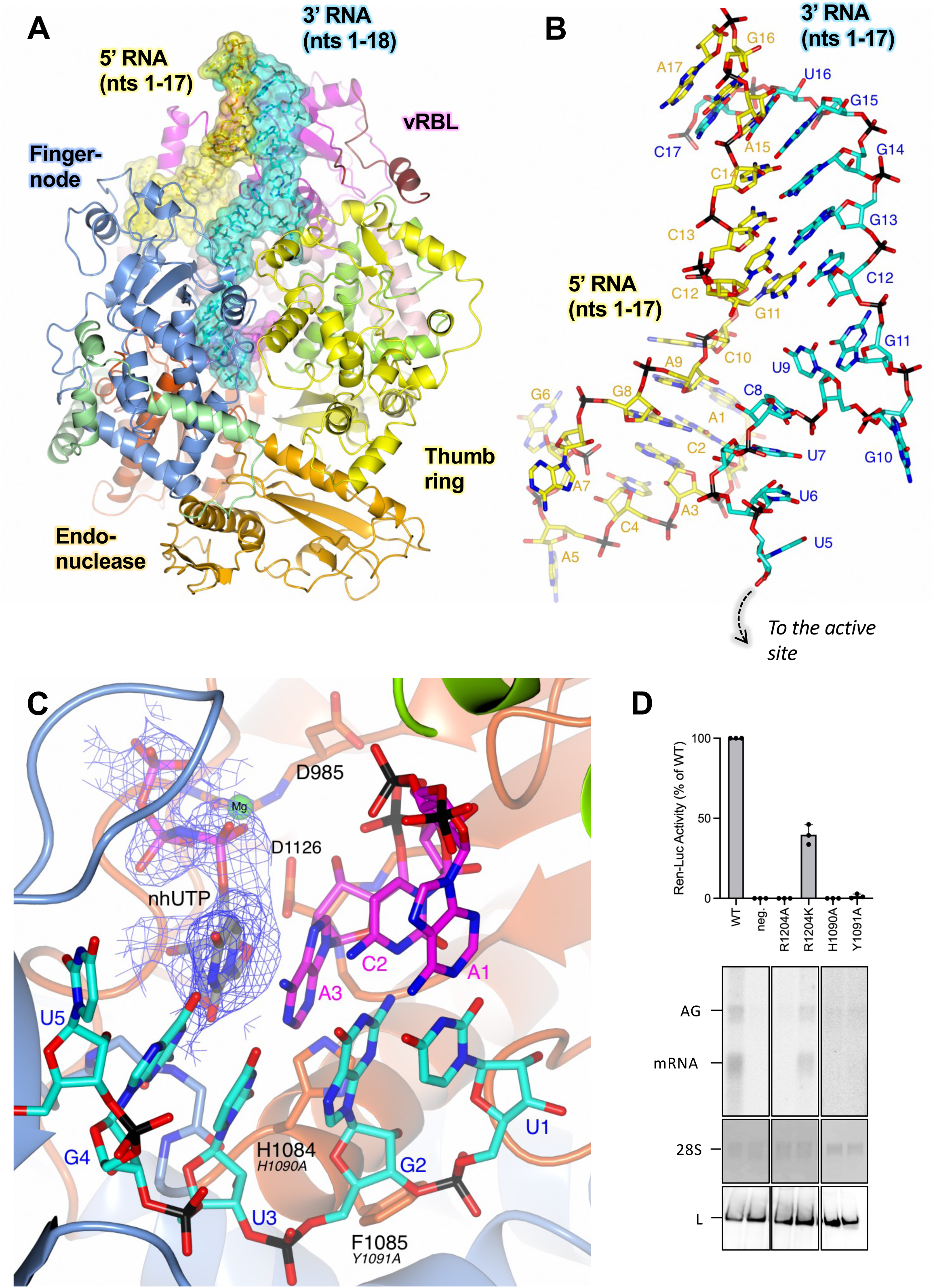
The L protein at early-elongation. (A) The SFTSV L protein (EARLY-ELONGATION) is shown with key L protein domains coloured according to the following colour scheme: endonuclease (orange), endonuclease linker (light green), PA-C like domain (light pink), vRBL (magenta), fingers and finger node (blue), thumb (green), and thumb ring (yellow). RNAs are shown as sticks with surface overlaid (50% transparency) and coloured either yellow (5′ cRNA), cyan (3′ cRNA), or magenta (progeny RNA). (B) A close-up of the 5′/3′ distal duplex in the EARLY-ELONGATION structure is shown with RNA nts labelled starting from either the 5′ end for the 5′ cRNA or from the 3′ end for the 3′ cRNA. (C) The active site in the EARLY-ELONGATION structure is shown with the EARLY-ELONGATION map overlaid. The map is clipped to the nhUTP and contoured to 4 σ. Residues from the SFTSV L protein are labelled and coloured according to the assigned domain (blue for the fingers domain, and dark orange for the palm domain) with the corresponding RVFV L protein residue shown in black italicised text underneath. (D) RVFV mini-replicon data for L protein with mutations in the vicinity of the active site presenting luciferase reporter activity (in standardised relative light units relative to the wild-type L protein (WT)). Data were presented as mean values ± SD of three biological replicates (n = 3). All biological replicates are shown as black dots (top panel). Middle panels present Northern blotting results with signals for antigenomic viral RNA (AG), viral mRNA (mRNA) and 28 S ribosomal RNA (28 S) as a loading control, and the bottom panel shows Western blot detection of FLAG-tagged L proteins (L) to demonstrate general expressability of the mutants. Uncropped original blots/gels are provided in the Supplementary Data.

In both the EARLY-ELONGATION and EARLY-ELONGATION-ENDO structures, we can reliably fit the 3′ cRNA residues found in the space between the distal duplex and the template entry channel. Notably, the G11 base stacks against the K533 sidechain whilst also interacting directly with the H535 sidechain (N). There is a sharp turn which allows the G10 base to sit in a pocket delineated by W1342 – K1347 and L1399 – S1400 (Supplementary Figure 12). While, the bases U9, C8, and U7 can be fit, this region is by comparison to the rest of the 3′ cRNA less defined in both maps, which is most likely due to a low number of interactions with protein residues. The C8 base interacts with the S561 sidechain and the U7 backbone phosphate is coordinated by the K1401 sidechain (Supplementary Figure 12). The remaining template RNA nts 1-6 of the 3′ cRNA, are overall much better defined and can be fit with high confidence in both structures. Bases U6 and U5 are found in the template entrance channel and G4 at the +1 position in the active site (Supplementary Figure 12). The only close interaction between the L protein and U6 and U5 is between the R871 sidechain and the U5 phosphate. In addition to the stalling nhUTP molecule, the G4 nt is coordinated by the Y923 and Q1080 carbonyls. There is only one close interaction between the remaining 3′ cRNA and the L protein and this is between the G1081 carbonyl and the U3 ribose.

The stalling nhUTP contributes towards the coordination of a single Mg^2+^ ion, also found at the +1 position, with each of the nhUTP phospho-oxygens contributing to one coordinating bond (Figure 3C). The A986 carbonyl and sidechains of D985 and D1126 provide the three remaining coordination bonds required to complete the bipyramidal Mg^2+^ coordination shell. The volume for the 3 nt product strand in both early elongation structures is noisy but nevertheless good enough to fit the cognate bases for the template U1, G2, and U3 bases, which are A1, C2, and A3, respectively (Supplementary Figure 13). There are some interactions between the L protein and product RNA nts but these are limited. A3 is stabilised by interactions from the S1125 and D1126 sidechains, which interact with the ribose O2’ and O3′ hydroxyls. There is also a relatively long interaction between the A3 phosphate and the S1183 sidechain. G2 is sandwiched in between A1 and A3 while the G2 phosphate interacts with the R1197 sidechain of L. There are no close protein interactions stabilising A1, which likely contributes to the poor map quality around this particular nt.

### The L protein undergoes significant remodelling as elongation progresses

There are several noticeable changes to the L protein comparing pre-initiation to early-stage and then late-stage elongation. The most apparent being an expansion of the L protein core, which leads to an increase in the internal diameter of the L protein by ~6 Å (Supplementary Figure 14). This opening up allows the L protein core to accommodate not just the small 3 nt product we observe in the EARLY-ELONGATION and EARLY-ELONGATION-ENDO structures but also the 9-bp product-template duplex we find in the LATE-ELONGATON structure. Additionally, in comparison to the apo-structure there is a rotation of the endonuclease domain by ~30° along the *Z* axis towards the L protein core (Figure 4A). This movement buries α4 of the endonuclease (residues 134 – 159, [24]) deeper within the L protein and, in doing so, partially opens the putative product exit channel in preparation for the later stages of elongation. It also provides space for the latter half of the thumb ring domain (roughly composed of L1509 – K1577) to rotate away from the L protein core, backfilling the space left free by the movement of the endonuclease domain (Figure 4A and B). Finally, a beta-hairpin of the thumb domain (T1333 – V1340) refolds and moves away from the L protein core (Figure 4B) unblocking the template exit channel.

**Figure 4.**
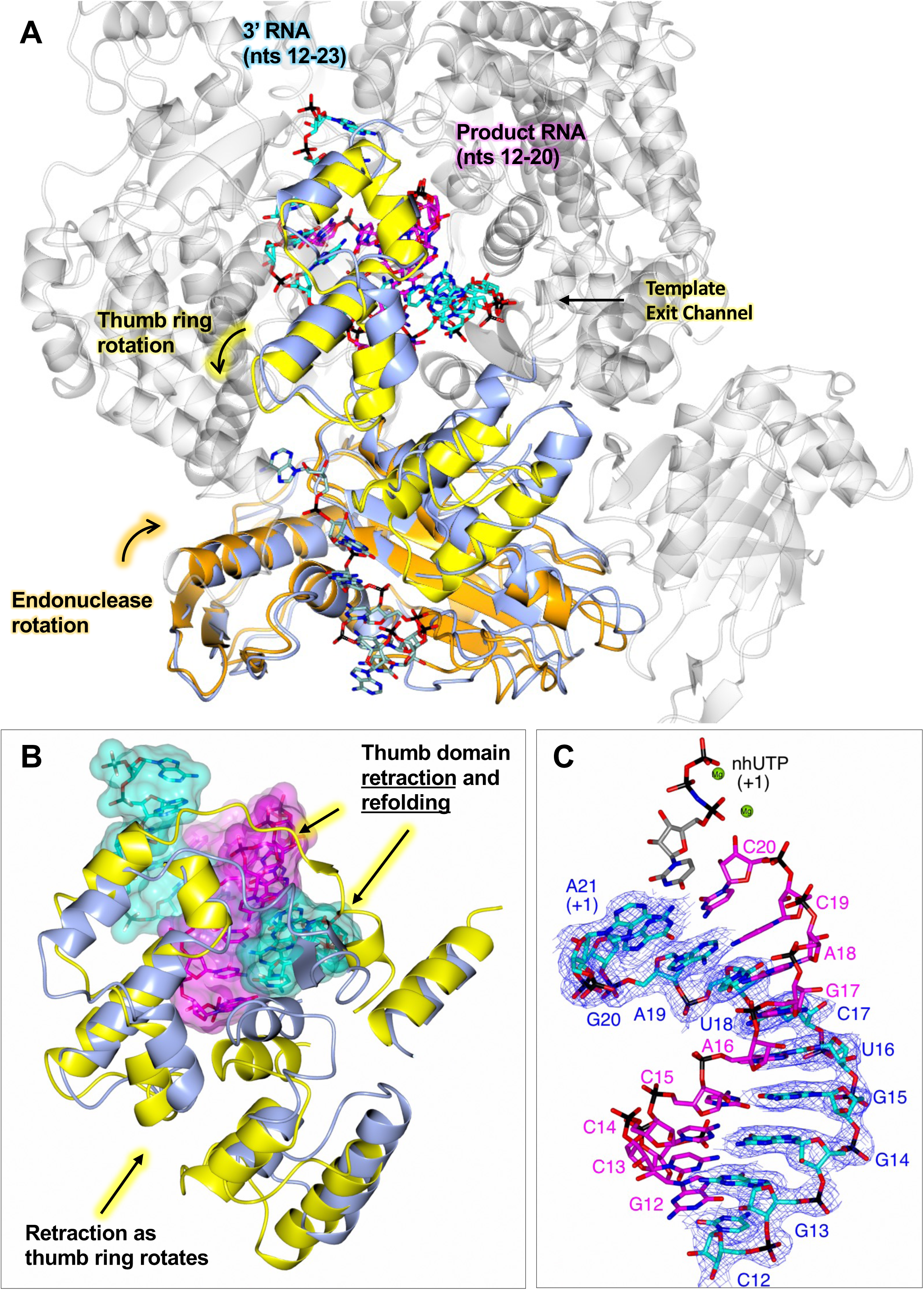
Conformational changes as the L protein proceeds from pre-initiation to late-elongation. (A) The SFTSV L protein (LATE-ELONGATION) is shown with the 5′ HOOK structure of the SFTSV L protein superposed by SSM in CCP4mg [51]. Parts in grey are very similar in both structures and only the LATE-ELONGATION structure is depicted. Domains of the LATE-ELONGATION structure that were shown to either rotate or remodel as the L protein moves from pre-initiation, i.e., hook binding, to late elongation are coloured (orange for the endonuclease and yellow for the thumb ring domain). The corresponding 5′ HOOK structure domains are shown in blue. RNAs are shown as sticks with surface overlaid (50% transparency) and coloured either cyan (3′ cRNA), magenta (progeny RNA), or pale green (endonuclease-bound RNA). Movements to the endonuclease and thumb ring domains precipitated by the progression to late elongation are indicated. (B) A close-up of the remodelling which leads to the opening of the template exit channel. As in A, the thumb domain and RNA of the LATE-ELONGATION structure is coloured yellow and cyan/magenta, respectively. The apo structure is shown in blue. (C) The 9-bp product-template duplex and the non-hydrolysable UTP (nhUTP) from the LATE-ELONGATION structure are shown as sticks with the LATE-ELONGATION map overlaid. The map is clipped to the 3′ cRNA strand and contoured to 4 σ. Catalytically important magnesium ions (Mg) are shown as green spheres.

### State of RNA at late-stage elongation

While in our early-stage elongation structures the 5′/3′ RNA distal duplex structure is maintained, in the later-stage LATE-ELONGATION structure, the distal duplex is disrupted completely. The SFTSV L protein is stalled through the incorporation of nhUTP at position A21 in the 3′ RNA template and the L protein inner cavity contains a 9-bp product-template duplex (Figure 4C). For related polymerases, such as LASV and LACV L proteins, the product-template duplex in the RdRp active site cavity was found to be 8 bp and 9-10 bp, respectively [13, 28]. The LATE-ELONGATION map is of sufficient quality and resolution (2.6 Å) to unambiguously identify each base in the growing strand. This led to the interesting finding that instead of stalling at the first A in the 3′ cRNA sequence (position 19), the L protein instead stalls at position 21 on the template, which is the second A in the overall sequence and the first A in the 6A tail of the modified template used.

### Product exit channel, template exit channel, and the 3′ secondary binding site

At the base of the SFTSV L protein core, the product-template duplex splits. This seems to be enabled partly by thumb-ring residue H1573, the sidechain of which protrudes directly into the path of the growing duplex. Unfortunately, the bottom of the SFTSV L protein core does not overlay nicely with the same space in the RVFV L protein. It was not possible therefore to explore the impact of mutating the equivalent of H1573 in the RVFV mini-replicon system. Additionally, recombinant expression of a D112A/H1573A SFTSV L protein mutant did not yield sufficient amounts of protein for *in vitro* testing. The growing strand then exits via the product exit channel, while the now used template exits via the template exit channel in the opposite direction. The putative product exit channel is open under apo/resting conditions and is formed of residues from the fingers (T839 and R835) and thumb ring domains (R1374, R1375, and R1526), as well as the endonuclease linker (E219 and E220) [30–33]. The sidechains of these residues create a pore through which the product RNA can leave the L protein core (Supplementary Figure 15).

The template exit channel is formed by residues from the thumb ring domains. In the published apo structures [30–33], this template exit channel is blocked. In the EARLY-ELONGATION and EARLY-ELONGATION-ENDO structures, we observe that the blocking motifs either rotate (L1509 – K1577) or refold (T1333 – V1340) and thereby, partially clear the channel aperture in preparation for template exit. As the L protein moves from early to late-stage elongation, the L protein core opens up and the associated global movement creates sufficient space for the template 3′ RNA to exit the L protein core via the template exit channel.

Several bunyaviruses and influenza viruses are known to coordinate 3′ RNA at secondary binding sites within the viral polymerase [13,28,29,53,55–57]. In the late-stage elongation structure reported here, we find nts 1-5 of the 3′ cRNA bound in a tunnel immediately adjacent to the vRBL (Figure 5A). This area broadly corresponds to the 3′ secondary binding sites observed in LASV [28] and LACV [13] L proteins. Unfortunately, in the polished LATE-ELONGATION map, it is not possible to fit an unbroken chain of RNA between the 3′ cRNA in the L protein core and the RNA bound to the 3′ secondary binding site. However, analysis of the surface electrostatics in this area reveals a positively charged cleft connecting the template exit channel to the 3′ secondary binding site (Figure 5B). We hypothesized that the lack of volume connecting the 3′ vRNA in the L protein core and the RNA bound to the 3′ secondary binding site might be due to over-sharpening of the map in this area.

**Figure 5.**
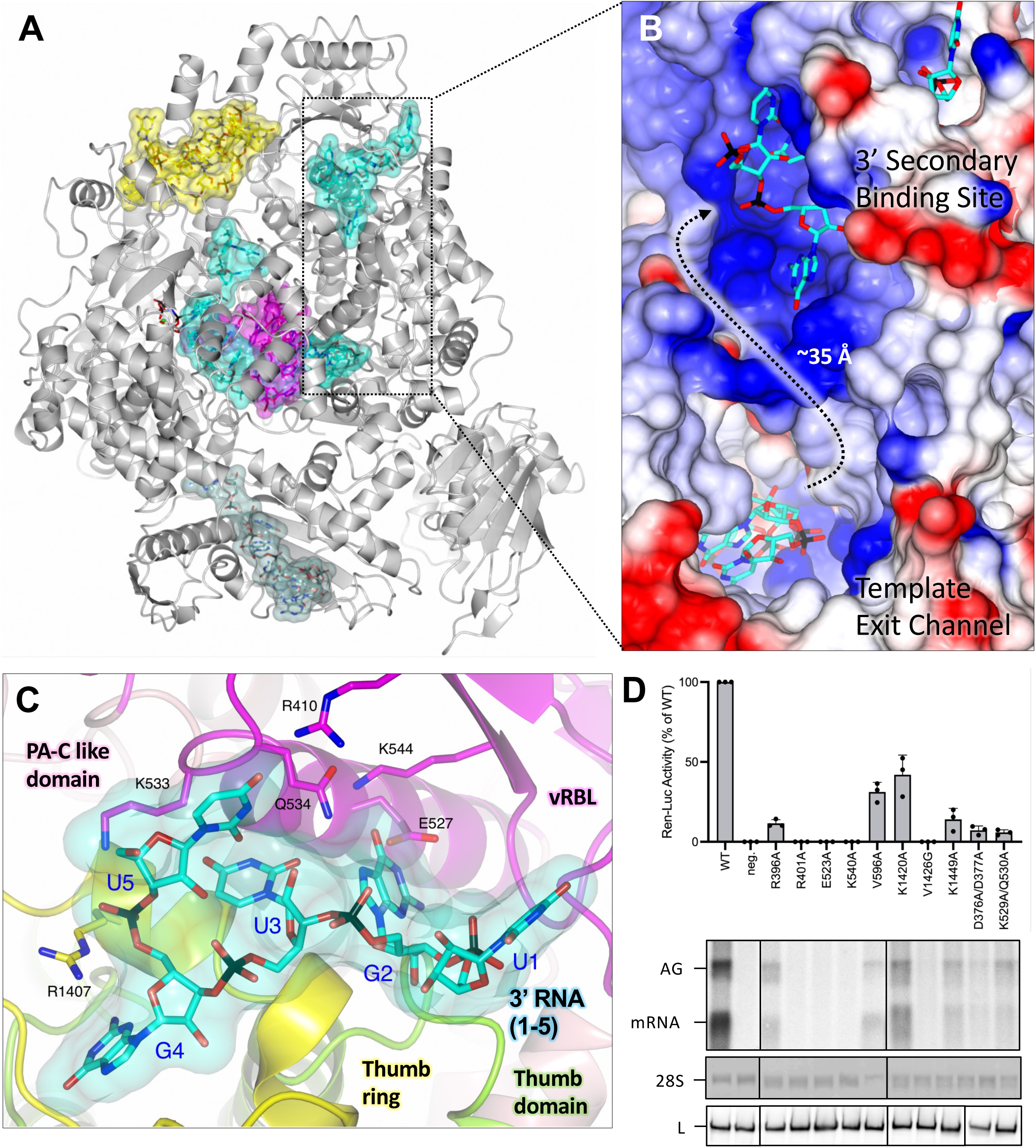
Interaction of 3′ cRNA at the 3′ secondary binding site. (A) The SFTSV L protein (LATE-ELONGATION) is shown with the protein depicted as ribbon in grey. RNAs are shown as sticks with surface overlaid (50% transparency) and coloured either yellow (5′ cRNA), cyan (3′ cRNA), magenta (progeny RNA), or pale green (endonuclease-bound RNA). (B) A close-up of the indicated area in A is shown as surface coloured by electrostatic potential demonstrating that the path connecting the template exit channel and 3′ secondary binding site has a largely positive charge. (C) The 3′ secondary binding site is shown with the L protein vRBL in magenta, the PA-C like domain in pink, thumb in green and thumb ring in yellow. The 3′ cRNA nts bound to the 3′ secondary binding site are shown as sticks with surface overlaid (50% transparency). Side chains of key interacting amino acids, coloured according to the protein domain, are shown as sticks and labelled accordingly. (D) RVFV mini-replicon data for L protein with mutations in the 3′ secondary binding site presenting luciferase reporter activity (in standardised relative light units relative to the wild-type L protein (WT)). Data are presented as mean values ± SD of three biological replicates (n = 3). All biological replicates are shown as black dots (top panel). Middle panels present Northern blotting results with signals for antigenomic viral RNA (AG), viral mRNA (mRNA) and 28 S ribosomal RNA (28 S) as a loading control, and the bottom panel shows Western blot detection of FLAG-tagged L proteins (L) to demonstrate general expressability of the mutants. Uncropped original blots/gels are provided in the Supplementary Data.

To assess this, we produced a blurred LATE-ELONGATION map and then re-examined this area of the L protein. In this blurred map, although the volume between the template exit channel and the 3′ secondary binding site is still rather amorphous, we clearly see the volume connecting the 3′ cRNA in the L protein core with the RNA bound to the 3′ secondary binding site (Supplementary Figure 16). Considering that the sequence of the RNA in the 3′ secondary binding site matches the first 5 nts of the 3′ cRNA used for this structure, and that the distance from the template exit channel to the secondary binding site (~35 Å) fits with the estimated length of 6 nts of RNA that are missing in the 3′ cRNA chain, we concluded that the RNA bound to the 3′ secondary binding site is likely the tail-end of the 3′ cRNA stalled in the L protein core. This follows similar observations in other sNSV polymerases, including influenza virus polymerase complex, where after being copied, the template 3′ end rebinds to the polymerase in a secondary binding site analogous to the secondary binding site described here [53].

There are extensive protein-RNA interactions at the 3′ secondary binding site which stabilise the bound RNA (Figure 5C). The first nt fit into the map at this site corresponds to U1 in the 3′ RNA sequence and is found on the extremity of the L protein surrounded by several flexible sidechains (D383, K386 and E387), however, none of these form close interactions with the U1 base. This is not too surprising as we observed something similar for the LASV L protein [28]. Nucleotide G2 sits in a pocket delineated by sidechains of Y408, E527, K544, L1254, P1256, I1344, and V1413. The E527, K544, and Q534 sidechains contribute to the coordination bonds and further stabilise the G2 base. The base of the third nt, U3, interacts with the Q534 sidechain while the U3 phosphate interacts with the R1418 sidechain. The base of G4 is coordinated by the carbonyl of Y1341 and the Y1414 sidechain. The base of the fifth and final fit nt, U5, is sandwiched between the K405 sidechain on one side and the K533-Q534 sidechains on the other. There is also a direct interaction between the R410 sidechain and the base O4’ hydroxyl.

### Mutational analysis of residues at the 3′ secondary binding site

To test for the functional relevance of the secondary 3′ RNA binding site, we first compared the LATE ELONGATION structure to the RVFV L and found that several of the key interacting residues are highly conserved (Supplementary Figure 17). We then mutated these key residues and tested 8 single and 2 double mutants in the RVFV mini-replicon system (Supplementary Table 3, Figure 5D). The majority of the mutations resulted in a loss of L protein activity of more than 90%, including: R401A (SFTSV: R410), E523A (SFTSV: E527), K540A (SFTSV: K544), V1426G (SFTSV: V1413), as well as the double-mutants D376A/D377A (SFTSV: S382/D383) and K529A/Q530A (SFTSV: K533/Q534). The remaining mutations resulted in a significantly reduced but not totally abolished L protein activity (between 10 and 42% activity remaining), including R396A (SFTSV: K405), V596A (SFTSV: K599), K1420A (SFTSV: R1407), and K1449A (SFTSV: K1435). This demonstrates that 3′ RNA binding to the secondary binding site is essential for both genome replication and transcription, as also demonstrated for LASV L protein [28].

### Initiation and priming of genome replication by the SFTSV L protein

To test how the SFTSV L protein could initiate replication, we incubated the L protein with 5′ cRNA, 3′ cRNA and a short 2 nt primer (AC) in the absence of any NTPs. Given that the AC primer could only base-pair with either one of the two UG repeats at the 3′ end of the 3′ cRNA, we had hoped this might allow us to visualise a priming loop (as seen in influenza virus) [58] or prime-and-realign loop (as reported for LACV) [13]. Processing of this dataset gave one major class, which we have named a RESTING state structure. Here, the 5′ cRNA hook is bound in the normal manner nestled betwixt the vRBL, PA-C like domain and fingers, while the 3′ cRNA is bound to the 3′ secondary binding site. Whereas in the early-elongation structures in which the 3′ RNA clearly feeds into the L protein core, in the RESTING structure, the distal duplex is absent. The RESTING structure is reminiscent therefore of the LASV 3END-CORE (PDB: 7OEA) [28] structure in which excess 3′ vRNA binds to the 3′ secondary binding site instead of entering the L protein core. Although we clearly see 3′ cRNA binding to the 3′ secondary binding site, we also found that we could fit 4 nts of the template 3′ cRNA in the L protein core, however, only 2 of these (U3 and G4) could be fit with confidence. The region of the map where these 3′ RNA nts are found is rather noisy but we think that similar to our early-elongation structures, the nts in the L protein core are likely the start of the 3′ cRNA. Although we could not visualize initiation, given that the 3′ cRNA repeatedly appears to reach at least 4 nts into the L protein core, and that we detect an RNA product corresponding exactly to the terminus of the 3′ template without additional or missing nts in our biochemical assays [17], we think that for vRNA synthesis SFTSV primes internally and then realigns this primer to the 3′ template extremity before proceeding with replication. How this works for transcription remains unclear and needs further investigation.

### Insights into the role and activity of the SFTSV L protein endonuclease

Inspection of our LATE-ELONGATION structure revealed, quite unexpectedly, RNA bound across the SFTSV L protein endonuclease (Supplementary Figure 18A). Analysis of the surface electrostatics in this area demonstrates that the RNA is bound across a positively charged cleft that includes the metal-coordination site and active site (Supplementary Figure 18B). Here, we can fit a ~7 nt RNA bound across the SFTSV L protein endonuclease domain. It is important to note that, in these experiments, the L protein used should be endonuclease inactive due to the mutation D112A in the active site [17, 24]. This is presumably why we observe uncleaved RNA. The volume for the endonuclease-bound RNA is of variable quality and it is difficult to determine base identity. However, we could fit the following sequence: 5′-ACACAGA-3′ and propose that the RNA bound to the endonuclease is likely an excess of 5′ cRNA. Indeed, in our cryo-EM experiments the 5′ cRNA is used at a 2-fold molar excess compared to the L protein.

There are no close interactions between the first two nts, A1 and C2, and the endonuclease or other parts of the L protein. The A3 phosphate forms an interaction with the R1532 sidechain of the thumb ring domain. The next two nts, C4 and A5, sit across two pockets which form the endonuclease metal-coordination site. In influenza virus endonuclease, these are known as the P1 and P0 pockets, respectively [59]. Interestingly, despite mutation of one of the metal-coordinators (D112A), the map suggests there is a single Mg^2+^ ion coordinated by E126 and the A5 phosphate. While there are no close interactions between the C4 and A5 bases, the C4 phosphate interacts with the sidechain of Y149, as well as the carbonyl, amide, and sidechain of T110. In addition to coordinating the bound Mg^2+^ ion, the A5 phosphate forms a long interaction with the E126 sidechain. The next nt, G6, sits in a pocket equivalent of the P1 pocket in influenza viruses [59]. The G6 base is then sandwiched between the K1558 sidechain and adjacent A5 base. The G6 phosphate meanwhile interacts with the amide and sidechain of T129. There is then a sharp turn which sees the base of the next nt A7 sandwiched between the sidechains of R131 and R832. There is also an interaction between the A7 phosphate and the R131 sidechain and between the A7 ribose ring-oxygen and the amide and sidechain of S132. At this point, the quality of the map degrades to such an extent that the fitting of further nts to determine the RNA sequence is no longer possible.

## DISCUSSION

The structural and functional data presented here provide insights into how the SFTSV L protein replicates the viral genome (Figure 6). First, in the pre-initiation state, compared to the apo conformation there is a relaxation of the vRBL and elements of the PA-C like domain revealing a positively charged cleft into which the 5′ RNA binds in a hook-like conformation. This is followed by the formation of a distal duplex structure between the 5′ and 3′ RNA which orients and positions the template RNA 3′ terminus such that it can feed into the L protein core and be positioned in the RdRp active site in preparation for initiation. Previously, we had proposed two alternative mechanisms for *de novo* initiation of genome replication by the SFTSV L protein [17]. Either the L protein initiates terminally at nt 1 of the template and then directly proceeds to elongation or the L protein initiates internally at position 3 of the template generating either an AC or ACA primer, which is then realigned to positions 1 and 2 or 1-3 of the template RNA before elongation [17]. In both cases, the end result is a blunt-end product and so it is difficult to discriminate between the two mechanisms biochemically. Furthermore, a prime-and-realign scenario has already been detected for several other sNSVs such as influenza viruses, peribunyaviruses and arenaviruses [13,15,60–64].

**Figure 6.**
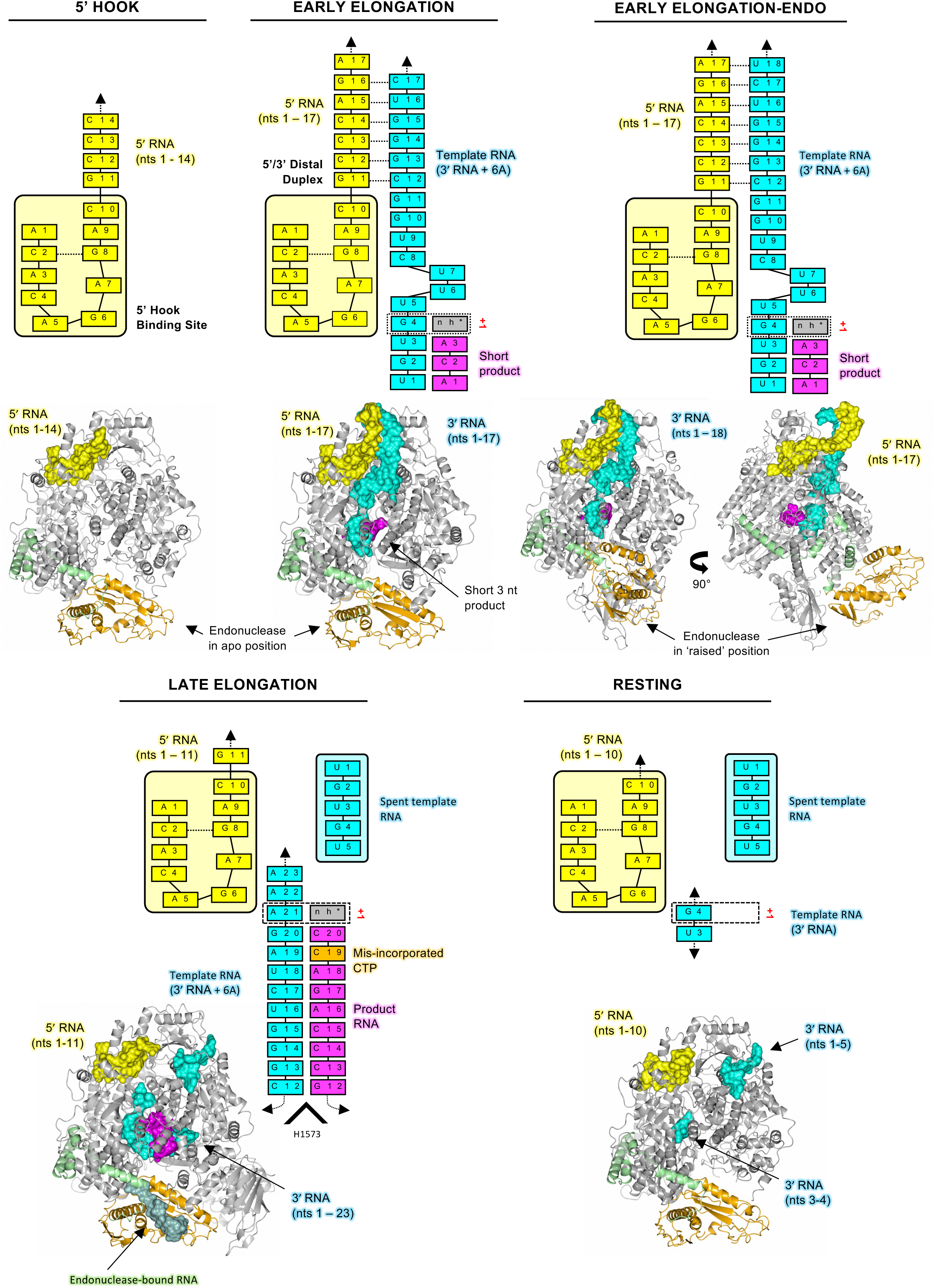
Overview of the SFTSV L protein at resting, pre-initiation, early-elongation, and late-elongation. A cartoon representation of each SFTSV L protein structure is shown in grey with the endonuclease and endonuclease-linker highlighted in orange and green, respectively. RNAs are shown as surface and coloured either yellow (5′ cRNA), cyan (3′ cRNA), magenta (progeny RNA), or pale green (endonuclease-bound RNA). In addition, the RNAs present in each structure are shown schematically and labelled.

Notably, in the early elongation structures, we found the 3′ cRNA positioned at least 4 nts into the RdRp active site suggesting an internal initiation scenario requiring subsequent realignment. Given the lack of an initiation stage structure, it is however not possible to reliably identify a potential priming or prime-and-realign loop which, in related RdRps, is stabilizing the first nucleotide and directing realignment. The prime-and-realign loop in LACV was suggested to work initially by stabilising the 3′ vRNA in a position which allows an AGU triplet to form at positions 4 to 6 on the 3′ vRNA [13]. By releasing the tension built up upon template progression, the prime-and-realign loop then supposedly pushes the 3′ vRNA backwards whilst the AGU triplet remains in the active site, thereby repositioning the AGU triplet to positions 1 to 3 on the 3′ vRNA [13]. Despite the lack of direct structural evidence, we made several observations that could be important for the SFTSV *de novo* initiation mechanism of genome replication. Firstly, we note that in our early-stage elongation structures, there is a bunching of the 3′ RNA at the template entrance channel with nts U6, U7, and C8 forming a kink. Secondly, G10 of the 3′ RNA is anchored in a pocket delineated by residues W1342 to K1347 as well as L1399 to S1400. Thirdly, the distal duplex stays intact even as the template reaches 4 nts into the active site. Finally, while the vast majority of amino acid residues in the L protein core overlay nearly perfectly in each of our structures (accounting for the general opening up of the inner cavity itself), we note that the sidechain of R1197 adopts one of two conformations. In fact, it is pointing either upward into the L protein core (pre-initiation) or laying flush with the growing RNA duplex (elongation). Although this difference is very subtle, the R1197 is of particular interest as it sits around 4-5 nts into the L protein core. Moreover, we know from our mutational analysis that mutating the equivalent of SFTSV L R1197 in the RVFV L protein (R1204) to a K reduces the L protein activity by over half and mutating it to an A completely abolishes L protein activity.

Based on the above, it seems possible that, at initiation, the R1197 sidechain hinders progression of the internally initiated product-template duplex while the distal duplex region of the promoter is still intact. Tension created in the template strand by the attempted template progression could lead to the template slipping backwards in the active site leading to the compressed RNA formation (nts 6 – 8) we see in our early elongation structures and thereby realignment of the primer to the 3′ terminus – in effect, a ‘*spring-loaded*’ prime-and-realign mechanism. This would result in the blunt-end product that we detect in our biochemical assays and also explain the lack of an obvious priming loop or prime-and-realign loop as seen in other polymerases. The blunt-ended template-product duplex might then be able to push the R1197 side chain aside upon strand progression without any further re-alignment. Of course, this proposed mechanism is speculative and the possibility remains that a potentially very flexible priming or prime-and-realign loop exists in a similar fashion to that seen in either influenza virus or LACV [13, 58].

As the L protein moves from early-elongation to late-stage elongation we find that the 5′/3′ distal duplex structure dissolves. At late-stage elongation, progeny RNA is released via a channel adjacent to the endonuclease domain, whereas spent template RNA is fed across the surface of the L protein in the direction of a 3′ secondary binding site. In the LATE-ELONGATION structure, the RNA fit in the 3′ secondary binding site is 5′-UGUGU-3′, which corresponds to the first 5 nts of the 3′ RNA. We expect that the 3′ RNA terminus binds to the 3′ secondary binding site with the remaining spent 3′ RNA then free to bulge out from the L protein in a similar fashion to that proposed for influenza virus [53]. In the cell, this RNA would presumably then be encapsidated by the viral N protein maintaining the viral ribonucleoprotein complex. However, structural data on this process is currently missing. As observed for LASV L protein [28], binding of the 3′ vRNA to the secondary binding site and 5′ hook binding is not mutually exclusive. Likewise, we could not detect any major changes in the 5′ hook binding site upon 3′ cRNA binding to the secondary binding site, leaving it unclear what triggers the release of the 5′ terminus at late elongation to achieve complete genome replication. Looking at the resting state structure it is conceivable that binding of the 3′ end to the secondary binding site and re-binding of the 5′ hook after passing through the RdRp active site supports the distal duplex formation, which then serves to guide the 3′ template terminus into the active site leading to the re-formation of a pre-initiation complex. However, further work looking at this recycling episode of the genome replication process is needed before any solid conclusions can be made.

Endonuclease domains have now been identified in many different bunyavirus L proteins and in each of these, the endonuclease is located at the N-terminus of the L protein [12,19,21–23,25–27,65,66]. While we can fit the SFTSV endonuclease into each of our structures published here, in the LATE-ELONGATION structure we can also visualise RNA bound across the endonuclease active site making it the first structure of a bunyaviral endonuclease with bound RNA substrate. This allows us to explore how the SFTSV endonuclease engages with substrate RNA. It has been hypothesized from biochemical studies that the phenuivirus endonuclease prefers purine-rich RNA species as substrates [21]. However, we could observe only one base-specific interaction between the hydroxyl group of Y76 and a C base right in the centre of the bound RNA. At this position the protein and RNA seem too close to accommodate a G base, whereas an A could in theory fit (Supplementary Figure 18C). In addition, Y76 appears to be stabilized in its position by contacts to neighbouring side chains which form part of the endonuclease-linker arm. All other protein-RNA contacts in the LATE-ELONGATION structure were solely formed between the protein and the RNA backbone. Therefore, while we cannot find structural evidence for a preference of purine-rich substrates, we do think that the LATE-ELONGATION structure suggests a disadvantage for G-rich RNA substrates. In the endonuclease assays described by Jones *et al.*, the GA-rich RNA substrate mainly consisted of A bases, which would not be contradictory to our observations [21]. Furthermore, as the length of the bound RNA in our structure is rather short (only 7 nts are fit), we cannot exclude that there may be further protein-RNA contacts with longer RNAs leading to the detected lower cleavage efficiency of pyrimidine-rich RNA substrates compared to purine-rich RNA substrates.

In addition to sequence specificity, regulation of the SFTSV endonuclease is also of particular interest not least because the endonuclease domain is thought to be central to the bunyaviruses ability to snatch capped-RNA fragments from host cells. For LASV, we found that the endonuclease is autoinhibited by two alternative mechanisms: either an inhibitory peptide or the C-terminal helix of the endonuclease domain blocks the endonuclease active site [28]. It has been proposed that the C-terminal α-helix of the SFTSV endonuclease (residues 211 – 233) could be responsible for controlling substrate access to the endonuclease active site (Supplementary Figure 19A) [24]. However, in each of our cryo-EM structures, we find that at around 2-turns into this particular α-helix (~L214), the helical arrangement unfolds, the protein chain bends by ~90° and subsequently forms the EN linker (Supplementary Figure 19B). The exception to this observation is the EARLY-ELONGATION-ENDO structure. In this structure the helix still unfolds around the same place after around 2-turns, but instead of veering away from the endonuclease active site, the strand proceeds for an additional 4-5 amino acids (Supplementary Figure 19C), which sit on top of the endonuclease active site. In a superposition of the endonuclease-bound RNA in the LATE-ELONGATION structure and the EARLY-ELONGATION-ENDO structure, the described 4-5 additional residues clash with the endonuclease-bound RNA (Supplementary Figure 19D). Thus, it would seem that in the raised endonuclease conformation access to the endonuclease active site is precluded. The EARLY-ELONGATION-ENDO structure therefore likely demonstrates one mechanism of endonuclease regulation in SFTSV, which is similar to what has been observed for LASV [28] and compatible with the mechanism proposed by Wang *et al.* [24]. However, as in LASV, there may be further mechanisms of inhibition in SFTSV L we have yet to discover.

Another interesting observation is the apparent ability of the SFTSV RdRp to misincorporate nucleotides when access to particular NTPs was restricted. A relatively low fidelity is a common feature of sNSV RNA polymerases but we also recognise that *in vitro* assay conditions may well produce artifacts. In cells, levels of all NTPs are not equal. For example, CTP and GTP are present at relatively low concentrations (≤ 0.5 mM) when compared to UTP (~1 mM) [67, 68]. The level of ATP is coupled to the metabolic status of the cell and is normally > 2 mM [67, 68]. The intracellular NTP concentrations are therefore even higher than those used in our *in vitro* polymerase assay (~ 0.2 mM) and may positively influence L protein fidelity. The mononegavirus respiratory syncytial virus (RSV) polymerase is capable of producing new RNA either *de novo* or via primer extension [69]. However, in *in vitro* biochemical assays, it was found that the ratio of *de novo* vs primer extension RNA synthesis is influenced by NTP concentration reinforcing this idea that both the concentration of NTPs and the ratio of individual NTPs are important for RNA synthesis [69]. Indeed, one suggested mechanism of action of the antiviral ribavirin is described to work by competitively inhibiting the cellular enzyme inosine 5′-monophosphate dehydrogenase, which ultimately leads to GTP depletion [70]. In addition, at high concentrations (≥100 μM) ribavirin has been shown to be incorporated as a GTP analogue into nascent viral RNA acting as a mutagen in following rounds of genome replication and transcription [71, 72]. Further factors known to influence the fidelity of polymerases are the availability and concentration of divalent metal ions with Mn^2+^ leading to higher rates of misincorporation compared to Mg^2+^ [73]. Our results on the misincorporation of CTP for UTP and *vice versa* under our assay conditions emphasize that NTP concentrations and relative ratios are important to consider for *in vitro* drug testing against NSV RdRp targets. Although out of scope for this study, we think it would be particularly interesting to investigate whether bunyaviral infections have an influence on the intracellular NTP pools as significant changes in dNTP levels have recently been observed in bacteria in response to bacteriophage infection [74]. Related to this topic, the low fidelity of the RdRp somewhat limits the confidence in the RNA product modelled in our LATE-ELONGATION structure. In the corresponding *in vitro* reaction, we detect two products, one most likely with a misincorporated CTP instead of UTP at position 19 of the product as we used nhUTP (>95% purity). Therefore, although we modelled the 21mer product with the described misincorporation, the cryo-EM data are most likely a mixture of both the 19mer and the 21mer. Interestingly, the mismatch does not result in a distortion of the RNA duplex, a scenario which has been described in the literature both for DNA and RNA suggesting that the rare tautomers giving rise to these mismatches are in fact relatively widespread in nucleic acids [75, 76].

To conclude, the biochemical and structural data presented here has provided us with key insights into how genome replication is catalysed by the SFTSV L protein. We have shown that there are specific changes to the L protein that occur upon 5′ RNA hook binding and that several L protein domains adapt further upon addition of 3′ RNA to accommodate the 5′/3′ RNA distal duplex. Our data show that as the L protein moves from pre-initiation to late-stage elongation, the L protein undergoes further remodelling with the inner cavity of the L protein opening up to accommodate the 9-bp protein-template duplex. Similar to other sNSV’s, we demonstrate that the SFTSV L protein has a functional 3′ secondary binding site, which is occupied under both resting conditions and also during the later stages of elongation. This structural data is accompanied by a comprehensive mutational analysis allowing us to identify key amino acid residues involved in binding RNA at the 5′ RNA hook binding site and 3′ secondary binding site. Altogether, this data provides an excellent foundation up on which further questions can be addressed including how the SFTSV L protein catalyses viral transcription.

## DATA AVAILABILITY

Coordinates and structure factors or map included in this paper have been deposited in the Worldwide Protein Data Bank (wwPDB) and the Electron Microscopy Data Bank (EMDB) with the following accession codes: SFTSV L protein bound to 5′ promoter RNA [5′ HOOK] EMD-15607 PDB-8AS6; SFTSV L protein early elongation structure [EARLY-ELONGATION] EMD-15608 PDB-8AS7; SFTSV L protein early elongation raised endonuclease structure [EARLY-ELONGATION-ENDO] EMD-15610 PDB-8ASB; SFTSV L protein late elongation structure [LATE-ELONGATION] EMD-15614 PDB-8ASD; SFTSV L protein resting structure [RESTING] EMD-15615 PDB-8ASG.

## FUNDING

We acknowledge funding for this collaborative project by the Leibniz Association’s Leibniz competition programme (grant K72/2017). Part of this work was performed at the Cryo-EM multi-user Facility at CSSB, headed by K.G. and supported by the UHH and DFG (grants INST 152/772-1, 774-1, 775-1 and 776-1). In the framework of this project, S.T. benefited from a travel grant from the Leibniz Institute for Virology; E.Q. was supported by an individual fellowship from the Alexander von Humboldt Foundation and a Klaus Tschira Boost Fund; M.R. received funding from the German Federal Ministry for Education and Research (grant 01KI2019).

## ACKNOWLEDGEMENTS

We want to thank Stephan Günther and Tomas Kouba for support and helpful discussions throughout the project. We thank Imre Berger for providing the DH10EMBaY *E. coli* and the team of the Eukaryotic Expression Facility (EEF) at European Molecular Biology Laboratory (EMBL) Grenoble for support and advice. We also acknowledge support by Carolin Seuring, Cornelia Cazey and Ulrike Laugks for access to the Cryo-EM multi-user facility at CSSB and providing time for sample preparation, screening, and data collection; Wolfgang Lugmayr for help and support in using the CSSB partition on the DESY computer cluster for cryo-EM data processing. We also want to thank Benoît Arragain for feedback on the manuscript.

## CONFLICT OF INTEREST DISCLOSURE

The authors certify that they have no affiliations with or involvement in any organization or entity with any financial or non-financial interest in the subject matter or materials discussed in this manuscript.

---

**Supplementary Table 2.** Cryo-EM data collection, processing, refinement, and validation statistics. This table provides the parameters and statistics for the data collection, processing, refinement, and structure validation of the cryo-EM structures. Refinement statistics were generated using the Phenix package.

|  | 5' HOOK | EARLY ELONGATION | EARLY ELONGATION ENDO | LATE ELONGATION | RESTING |
| --- | --- | --- | --- | --- | --- |
| PDB Code | 8AS6 | 8AS7 | 8ASB | 8ASD | 8ASG |
| EMBD Code | EMD-15607 | EMD-15608 | EMD-15610 | EMD-15614 | EMD-15615 |
| Data collection and processing |  |  |  |  |  |
| Microscope | Titan Krios G3 | Titan Krios G3 |  |  | Titan Krios G3 |
| Voltage (kV) | 300 | 300 |  |  | 300 |
| Camera | Gatan K3 BioQuantum | Gatan K3 BioQuantum |  |  | Gatan K3 BioQuantum |
| Magnification | 105,000 | 105,000 |  |  | 105,000 |
| Nominal negative defocus range (μm) | 0.8 – 2.5 | 0.8 – 2.5 |  |  | 0.8 – 2.5 |
| Exposure time (s) | 2.5 | 2.5 |  |  | 2.5 |
| Electron exposure (e-1/Å²) | 55 | 53 |  |  | 55 |
| Frames collected (no.) | 50 | 50 |  |  | 50 |
| Frames processed (no.) | 50 | 50 |  |  | 50 |
| Pixel size (Å) | 0.85 | 0.85 |  |  | 0.85 |
| Micrographs (no.) | 3,178 | 3,547 |  |  | 2,985 |
| Total particle images | 1,648k | 2,651k |  |  | 278k |
| Refinement |  |  |  |  |  |
| Final particle subset (no.) | 60k | 89k | 16k | 60k | 37k |
| Map resolution (Å), 0.143 FSC | 3.4 | 2.6 | 3.6 | 2.6 | 3.2 |
| Map sharpening *B* factor (Å²) | -93 | -40 | DeepEMhancer | -37 | Local Filter |
| Map vs. model cross-correlation CC-MASK | 0.83 | 0.82 | 0.73 | 0.84 | 0.82 |
| Model composition |  |  |  |  |  |
| Non-hydrogen atoms | 12784 | 13431 | 14487 | 15533 | 12689 |
| Protein residues | 1566 | 1575 | 1715 | 1826 | 1544 |
| Nucleotide residues | 14 | 37 | 38 | 44 | 17 |
| Waters | 0 | 10 | 0 | 32 | 0 |
| Other | N/A | 2 Mg, 1 nhUTP, 1 HEPES | 1 Mg, 1 nhUTP | 3 Mg, 1 nhUTP | 1 Mg |
| **Mean B factors (Å)** |  |  |  |  |  |
| Protein | 66.16 | 42.93 | 78.81 | 41.97 | 68.49 |
| Nucleotide | 73.61 | 62.80 | 86.61 | 41.04 | 102.60 |
| Ligand | N/A | 47.44 | 71.60 | 27.38 | 47.52 |
| Water | N/A | 28.40 | N/A | 28.35 | N/A |
| **R.m.s. deviations** |  |  |  |  |  |
| Bond lengths (Å) | 0.004 | 0.003 | 0.003 | 0.004 | 0.003 |
| Bond angles (°) | 0.709 | 0.619 | 0.682 | 0.653 | 0.736 |
| **Validation** |  |  |  |  |  |
| MolProbity score | 1.70 | 1.47 | 1.73 | 1.42 | 1.70 |
| All-atom clashscore | 4.32 | 5.32 | 6.72 | 3.96 | 6.56 |
| Poor rotamers (%) | 0.00 | 0.29 | 0.07 | 0.06 | 0.29 |
| **Ramachandran plot** |  |  |  |  |  |
| Favoured (%) | 91.95 | 96.87 | 94.74 | 96.37 | 95.15 |
| Allowed (%) | 8.05 | 3.13 | 5.26 | 3.63 | 4.85 |
| Outliers %) | 0.00 | 0.00 | 0.00 | 0.00 | 0.00 |

**Supplementary Table 3.** Raw data from RVFV mini replicon assays. For each mutant, the calculated Ren-Luc activity (as a% compared to the WT) from 3 biological replicates (as indicated in the table), the calculated average and standard deviation is provided.

| RVFV Residue(s) | Replicate 1 | Replicate 2 | Replicate 3 | Average | Standard Deviation |
| --- | --- | --- | --- | --- | --- |
| **Set 1 – Hook Interacting Residues** |  |  |  |  |  |
| E312A | 101,18 | 102,42 | 123,95 | 109,18 | 10,45 |
| D313A | 50,48 | 113,73 | 100,23 | 88,15 | 27,20 |
| Q326A | 97,83 | 115,05 | 93,20 | 102,02 | 9,40 |
| G418V | 0,02 | 0,01 | 0,01 | 0,01 | 0,01 |
| K422A | 38,34 | 48,68 | 45,85 | 44,29 | 4,36 |
| R434A | 51,86 | 78,88 | 74,13 | 68,29 | 11,78 |
| K533A | 86,89 | 130,18 | 151,11 | 122,73 | 26,74 |
| F561A | 2,82 | 7,75 | 8,33 | 6,30 | 2,47 |
| K602A | 74,50 | 96,79 | 121,88 | 97,72 | 19,35 |
| K661A | 0,12 | 0,08 | 0,41 | 0,20 | 0,15 |
| R700A | 0,02 | 0,01 | 0,01 | 0,01 | 0,01 |
| V698G | 30,42 | 89,70 | 79,89 | 66,67 | 25,94 |
| E769A | 102,07 | 129,69 | 171,22 | 134,33 | 28,42 |
| S771A | 103,28 | 184,97 | 173,16 | 153,80 | 36,05 |
| I1041G | 4,38 | 5,65 | 8,13 | 6,05 | 1,56 |
| H1045A | 3,50 | 3,13 | 2,87 | 3,17 | 0,26 |
| V1055G | 42,86 | 82,81 | 48,38 | 58,02 | 17,68 |
| F1059A | 0,02 | 0,04 | 0,01 | 0,02 | 0,01 |
| S437A/K438A | 22,75 | 38,12 | 27,38 | 29,42 | 6,44 |
| D1049A/R1051A | 35,39 | 42,69 | 27,07 | 35,05 | 6,38 |
| **Set 2 – 3' Secondary Binding Site** |  |  |  |  |  |
| R396A | 10,17 | 14,25 | 10,17 | 11,53 | 1,92 |
| R401A | 0,03 | 0,03 | 0,07 | 0,04 | 0,02 |
| E523A | 0,02 | 0,01 | 0,01 | 0,01 | 0,01 |
| K540A | 0,01 | 0,01 | 0,02 | 0,01 | 0,00 |
| V596A | 36,44 | 32,44 | 24,65 | 31,18 | 4,89 |
| K1420A | 28,30 | 52,39 | 45,12 | 41,94 | 10,09 |
| V1426G | 0,07 | 0,01 | 0,03 | 0,04 | 0,02 |
| K1449A | 6,54 | 15,53 | 20,06 | 14,04 | 5,62 |
| D376A_D377A | 4,24 | 9,62 | 7,83 | 7,23 | 2,24 |
| K529A_Q530A | 5,89 | 7,56 | 4,17 | 5,87 | 1,39 |
| **Set 3 – Priming Loop-like/Active Site Residues** |  |  |  |  |  |
| R1204A | 0,04 | 0,08 | 0,03 | 0,05 | 0,02 |
| R1204K | 39,40 | 46,17 | 33,70 | 39,76 | 5,10 |
| H1090A | 0,03 | 0,03 | 0,01 | 0,02 | 0,01 |
| Y1091A | 0,95 | 0,21 | 2,90 | 1,35 | 1,14 |

**Supplementary Figure 1.**
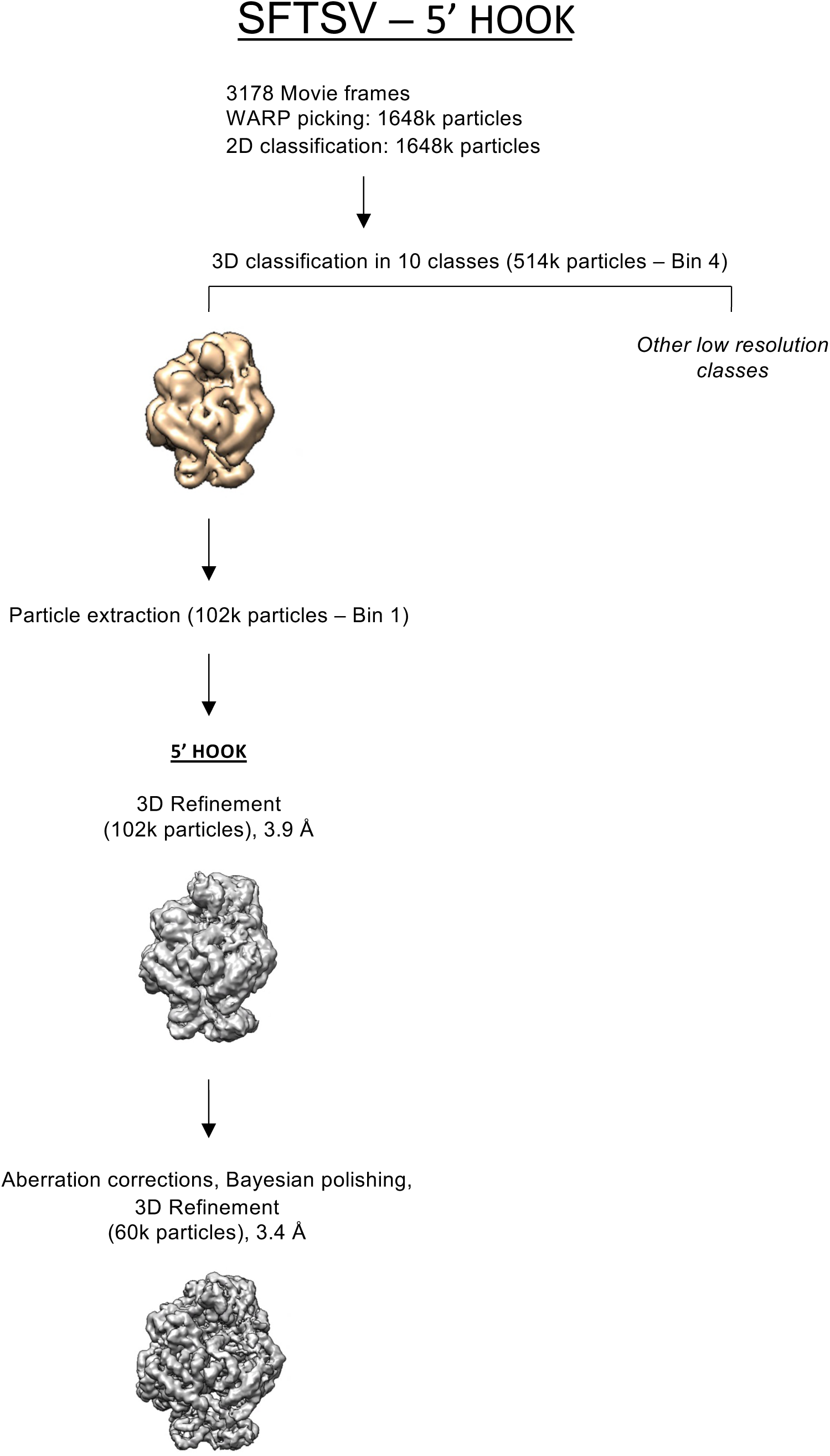
Cryo-EM data processing workflow for the SFTSV L 5’ HOOK dataset. After micrograph pre-processing and BoxNet particle picking in Warp [40], particles were extracted using a binning factor of 4 in RELION [37, 38] for 2D classification. 2D classes that had visible protein secondary structure were selected for subsequent 3D classification (514k particles). The particles were classified into 10 classes for 3D classification with angular assignment. Incomplete, low resolution or classes containing damaged particles were excluded from further data analyses. Particles from a single class, denoted 5’ HOOK, were re-extracted with the original pixel size value and refined before further 3D classification without angular assignment. The final particle subset was CTF refined, Bayesian polished and post-processed in RELION 3.1/4.0 [37, 38].

**Supplementary Figure 2.**
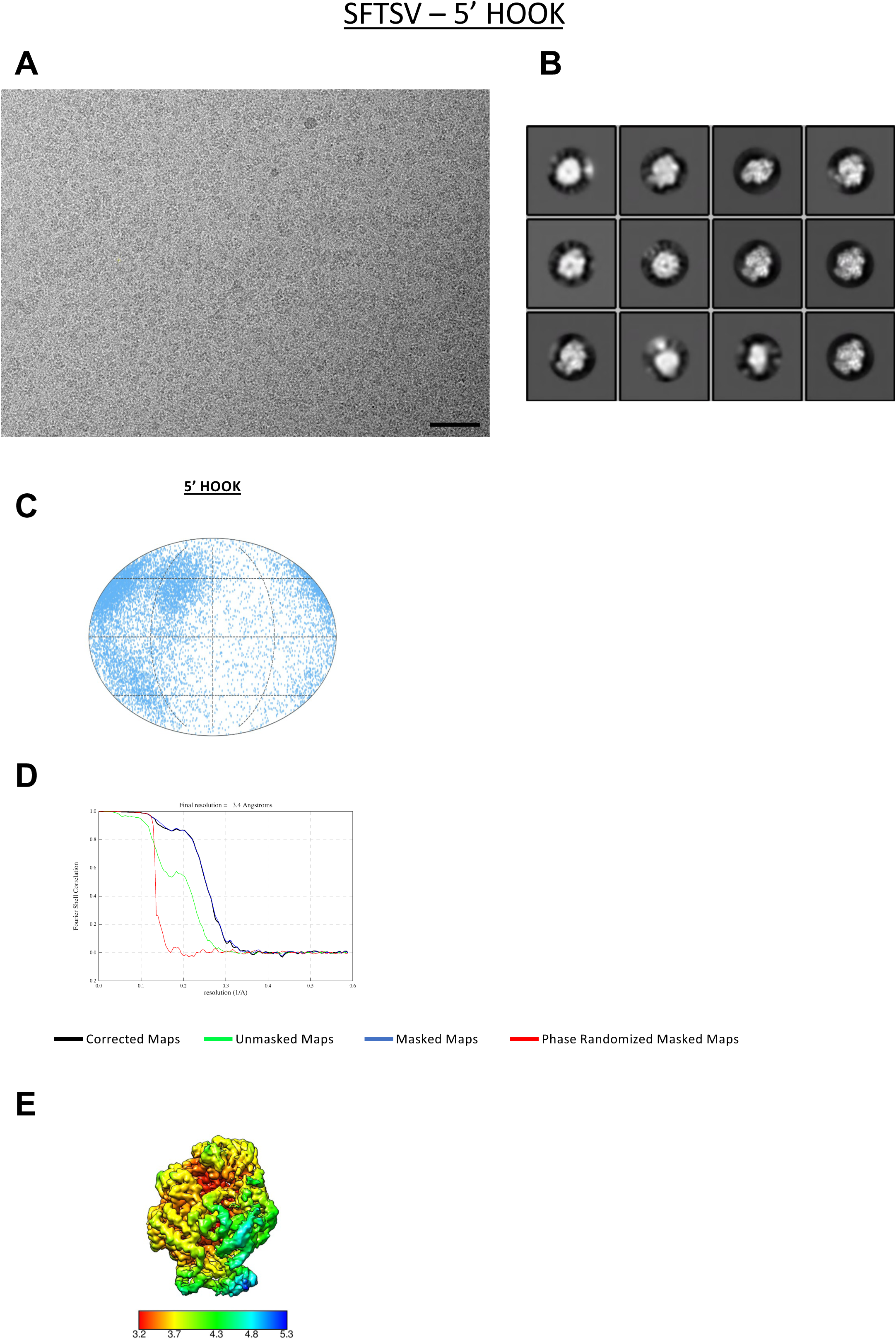
Cryo-EM of SFTSV L protein bound to the 5’ RNA (5’ HOOK). (A) A representative micrograph of SFTSV L protein in free standing ice after MotionCor2 [39] correction at defocus of −2.0 m. (B) Most populated 20 class averages of the of SFTSV L protein bound to the 5’ hook primer (see Materials and Methods). (C). Angular distribution for all of the particles refined visualized on a globe-like plane. (D) Fourier shell correlation (FSC) curves for the final refined class. The plot of the FSC between two independently refined half maps shows the overall resolution as indicated by the gold standard criteria of the FSC cut-off at 0.143. (E) Surface representation of the local resolution distribution of the final class after refinement. Maps are coloured according to the local resolution calculated within the RELION software package [37, 38] and displayed in Chimera [42]. Resolution is as indicated in the colour bar.

**Supplementary Figure 3.**
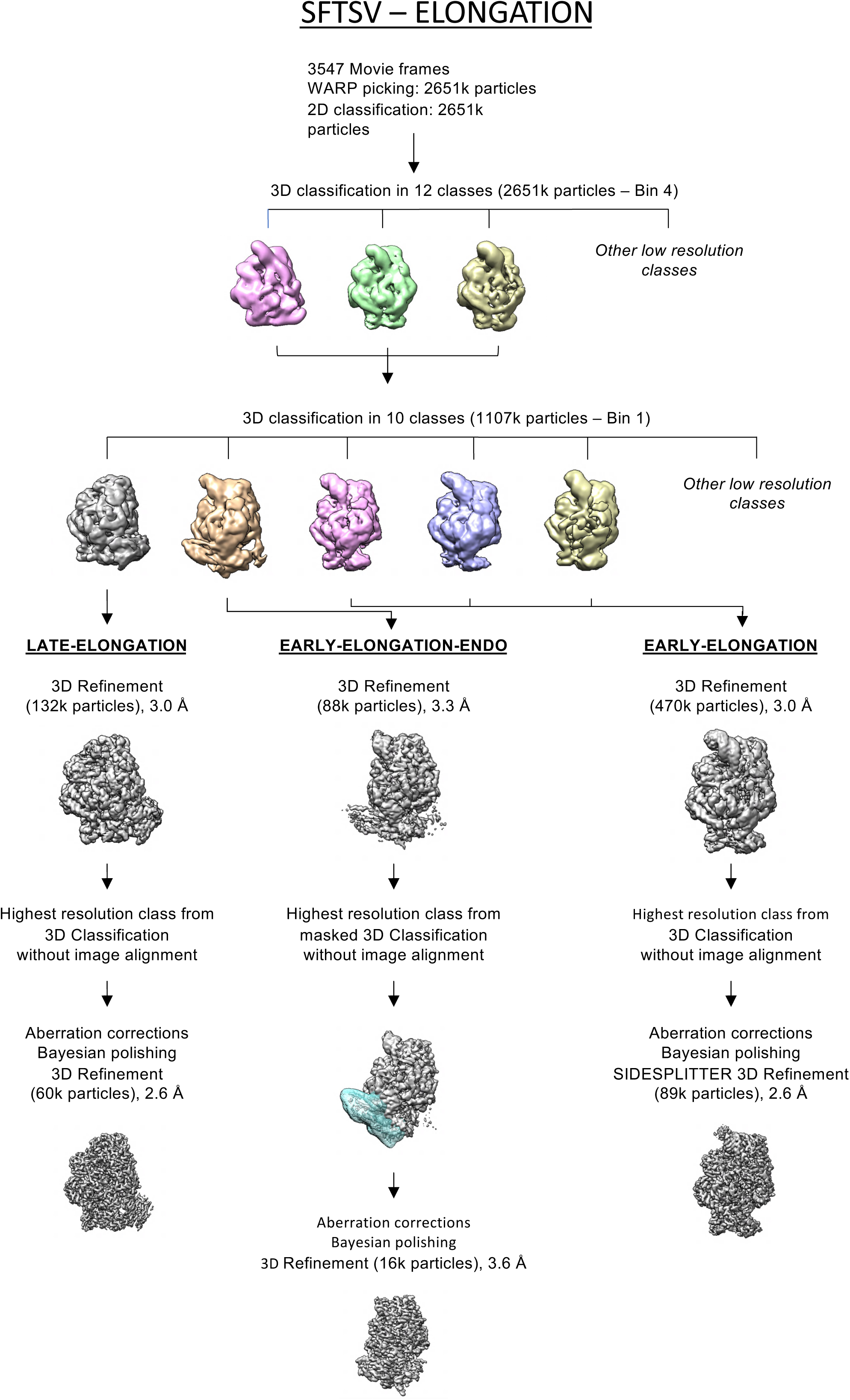
Cryo-EM data processing workflow for the SFTSV L elongation dataset. After micrograph pre-processing and BoxNet particle picking in Warp [40], particles were extracted using a binning factor of 4 in RELION [37, 38] for 2D classification. 2D classes were used for initial check but not to filter out particles so all picked particles went into 3D classification (2651k particles). The particles were classified into 12 classes for 3D classification with angular assignment. Incomplete, low resolution or classes containing damaged particles were excluded from further data analyses. The remaining particles were then re-extracted with original pixel size for further 3D classification using 10 classes. Among these, 3 classes with visible protein secondary structure were pooled as “EARLY ELONGATION”. These had density where we expect binding of an RNA duplex and were globally 3D refined. In parallel, 2 additional classes, each with distinct features, were refined independently. The so-called “EARLY ELONGATION ENDO” had a poor density for the endonuclease domain which was then masked for a subsequent 3D classification without image alignment. Afterwards, the highest resolution class showing a clear density for the endonuclease after refinement. The “LATE ELONGATION” class, was 3D refined before being further 3D classified without image alignment. The highest resolution class was then selected for further refinement. Final particle subsets were CTF refined, Bayesian polished and post-processed in RELION 3.1/4.0 [37, 38].

**Supplementary Figure 4.**
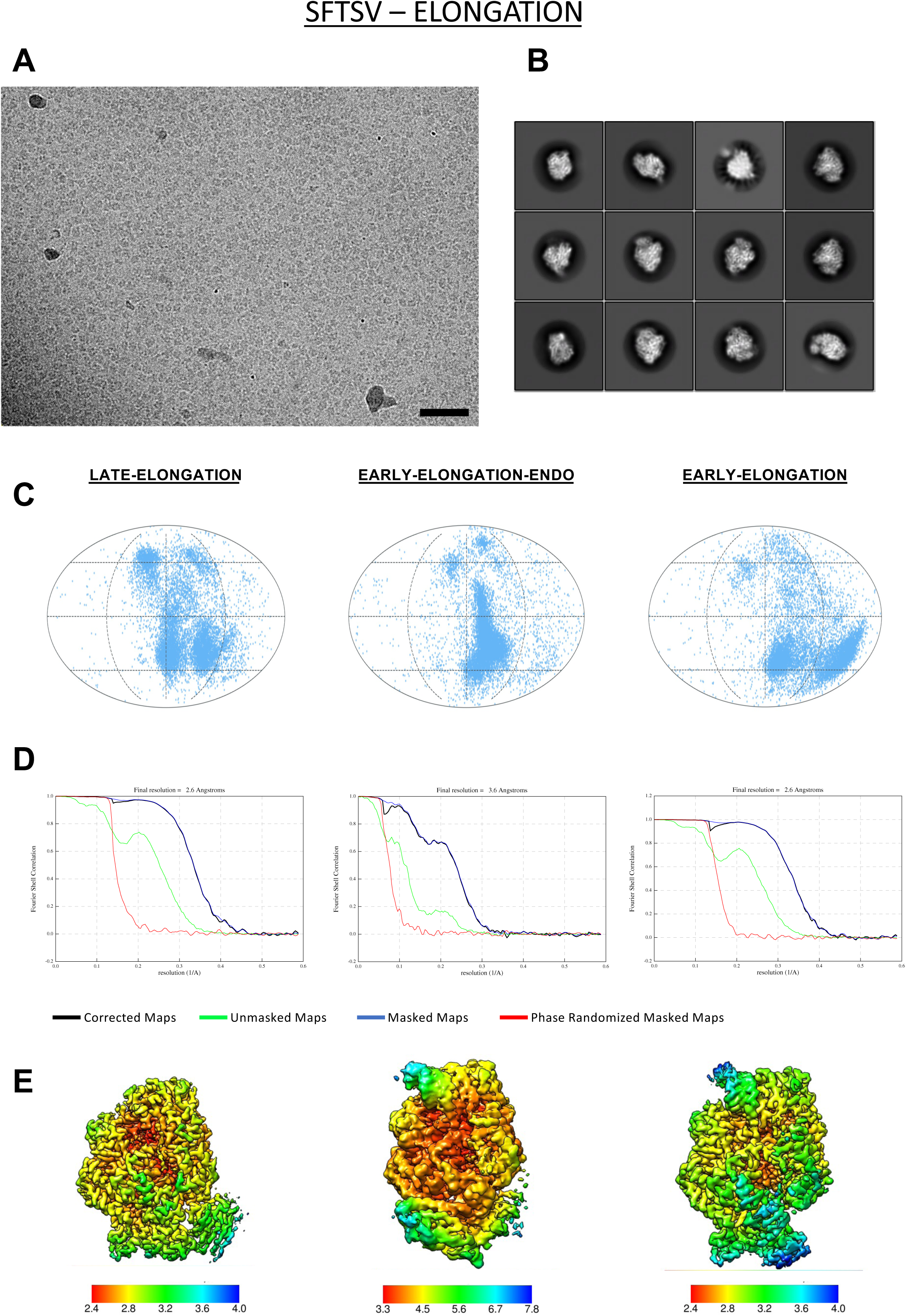
Cryo-EM of the SFTSV L protein elongation reaction. (A) A representative micrograph of SFTSV L protein in free standing ice after MotionCor2 [39] correction at defocus of −2.0 m. (B) A selection of the most populated 20 class averages of the of SFTSV L protein stalled with nhUTP (see Materials and Methods). (C) Angular distribution for all of the particles contributing to the different refined classes visualized on a globe-like plane. (D) Fourier shell correlation (FSC) curves for each final refined class. The plot of the FSC between two independently refined half maps shows the overall resolution as indicated by the gold standard criteria of the FSC cut-off at 0.143. (E) Surface representation of the local resolution distribution of each final class after refinement. Maps are coloured according to the local resolution calculated within the RELION software package [37, 38] and displayed in Chimera [42]. Resolution is as indicated in the colour bar.

**Supplementary Figure 5.**
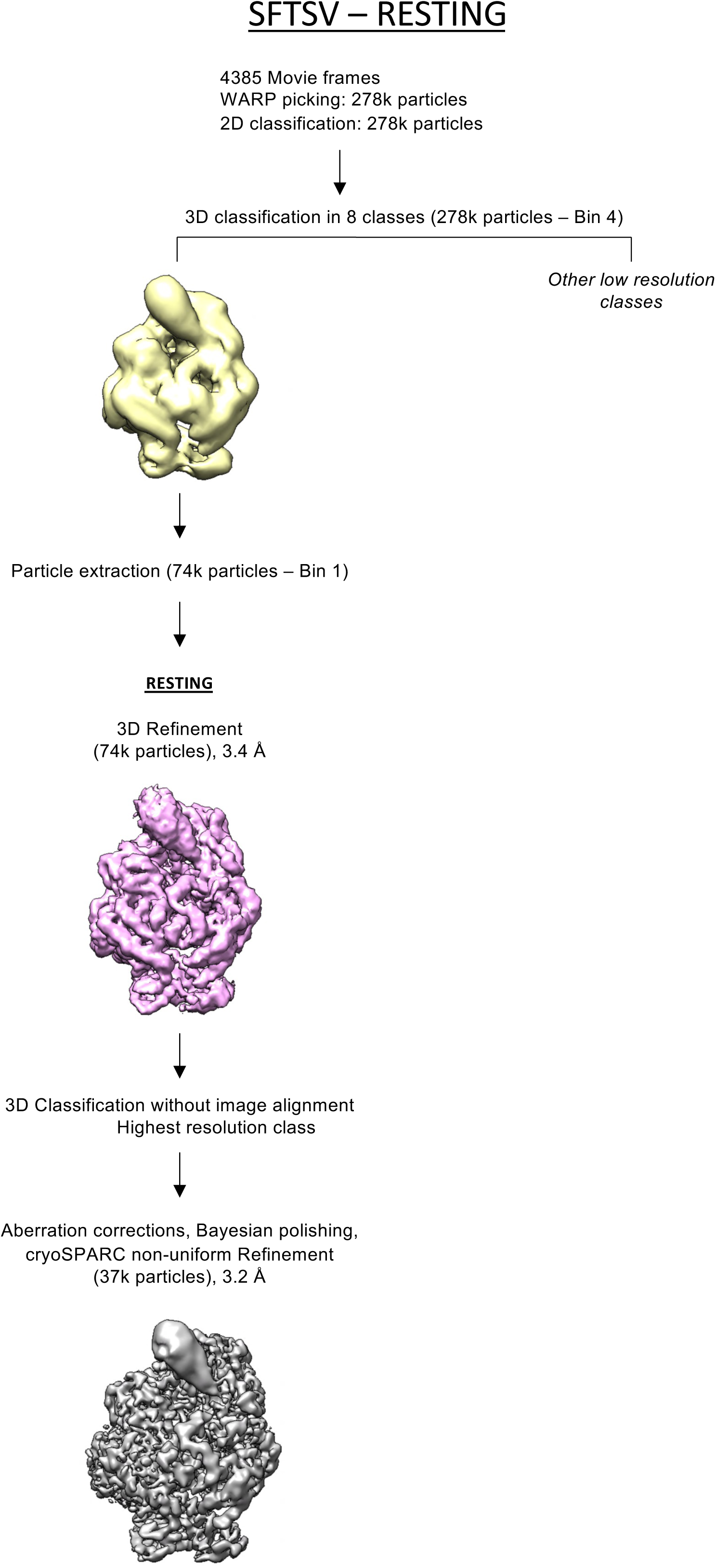
Cryo-EM data processing workflow for the SFTSV L RESTING dataset. After micrograph pre-processing and BoxNet particle picking in Warp [40], particles were extracted using a binning factor of 4 in RELION [37, 38] for 2D classification. 2D classes were used for initial check but not to filter out particles so all picked particles went into 3D classification (278k particles). The particles were classified into 8 classes for 3D classification with angular assignment. Incomplete, low resolution or classes containing damaged particles were excluded from further data analyses. A single class, “RESTING”, showed secondary structure features which was selected for further processing. Particles were re-extracted with the original pixel size value and refined before a subsequent 3D classification without image alignment with the highest resolution class selected for further 3D refinement. Final particle subset was CTF refined and Bayesian polished in RELION 3.1/4.0 [37, 38] prior to being subjected to non-uniform refinement and post-processed in cryoSPARC [43, 44].

**Supplementary Figure 6.**
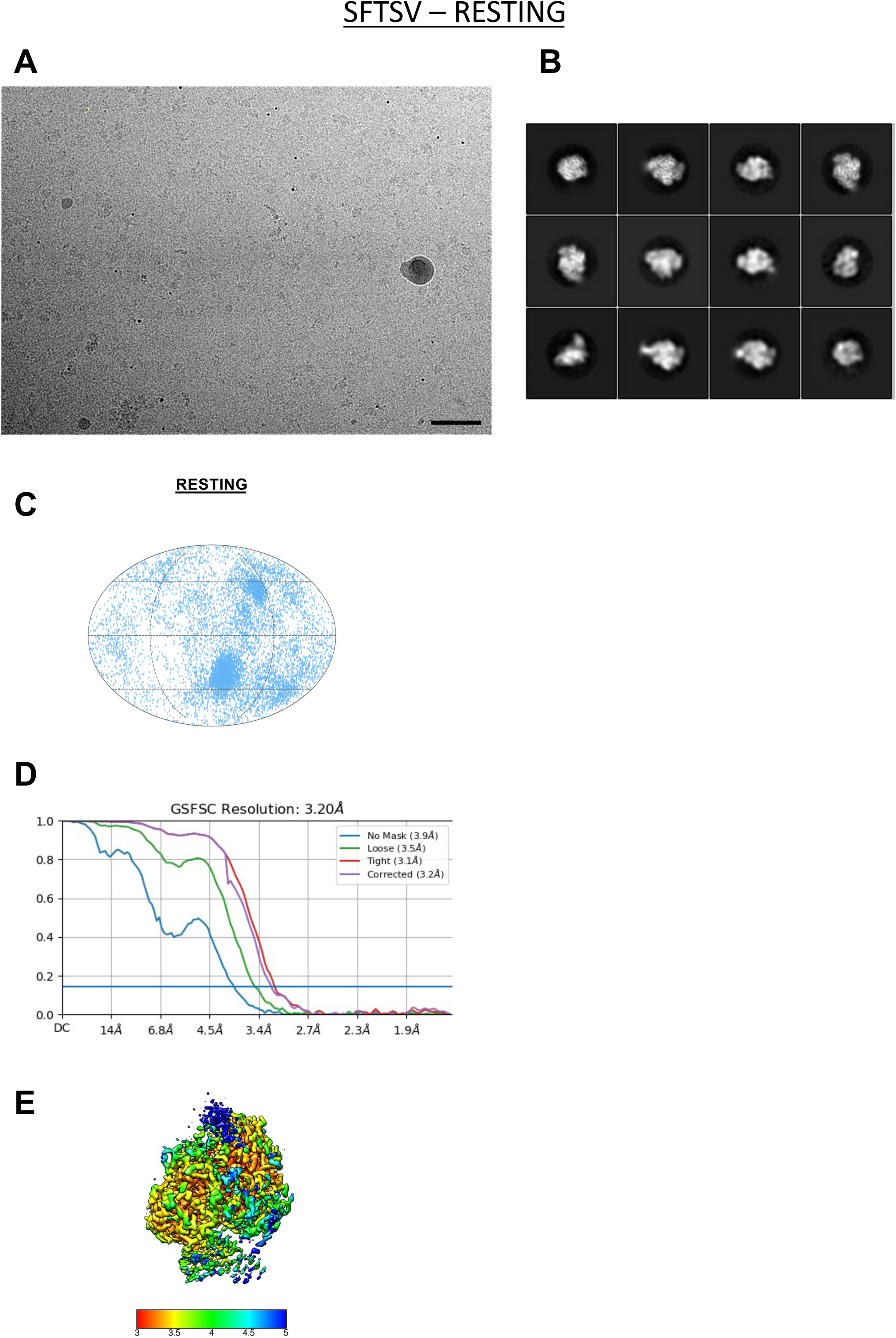
Cryo-EM of the SFTSV L RESTING dataset. (A) A representative micrograph of SFTSV L protein in free standing ice after MotionCor2 [39] correction at defocus of −2.0 m. (B) A selection of the most populated 20 class averages of the SFTSV L protein bound to AC primer (see Materials and Methods). (C) Angular distribution for all of the particles refined visualized on a globe-like plane. (D) Fourier shell correlation (FSC) curves for the final refined class. The plot of the FSC between two independently refined half maps shows the overall resolution as indicated by the gold standard criteria of the FSC cut-off at 0.143. (E) Surface representation of the local resolution distribution of each final class after refinement. Maps are coloured according to the local resolution calculated within the RELION software package [37, 38] and displayed in Chimera [42]. Resolution is as indicated in the colour bar.

**Supplementary Figure 7.**
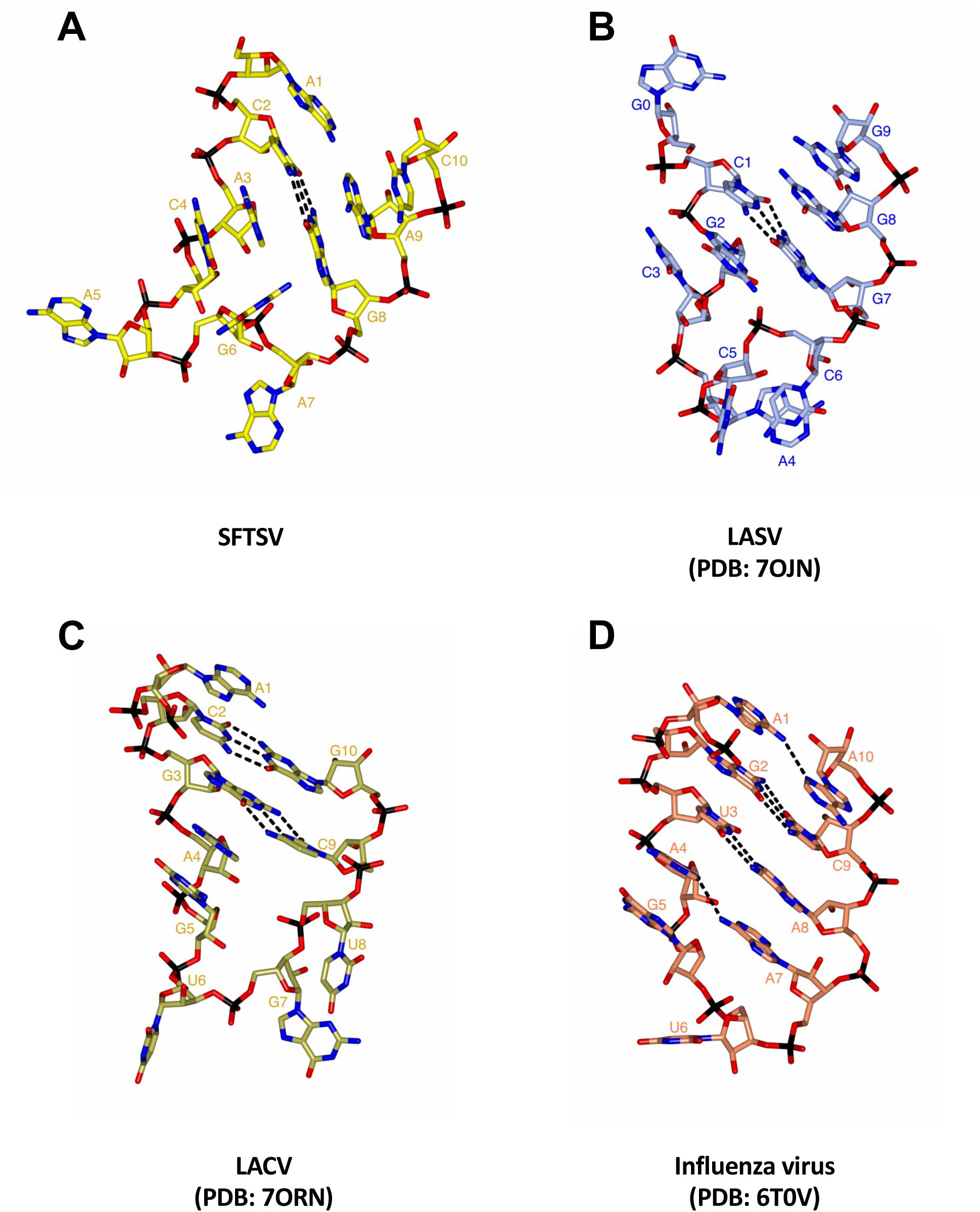
Comparison of known 5’ RNA hook structures for sNSVs. The 5’ RNA hook structures of SFTSV (A), LASV (B, nts 1-10 from PDB: ?OJN), LACV (C, nts 1-10 from PDB: ?ORN), and Influenza virus (D, nts 1-10 from PDB: 6T0V) are shown as sticks. Individual bases are labelled from the 5’ end and base-pairing interactions are shown as dotted lines.

**Supplementary Figure 8.**
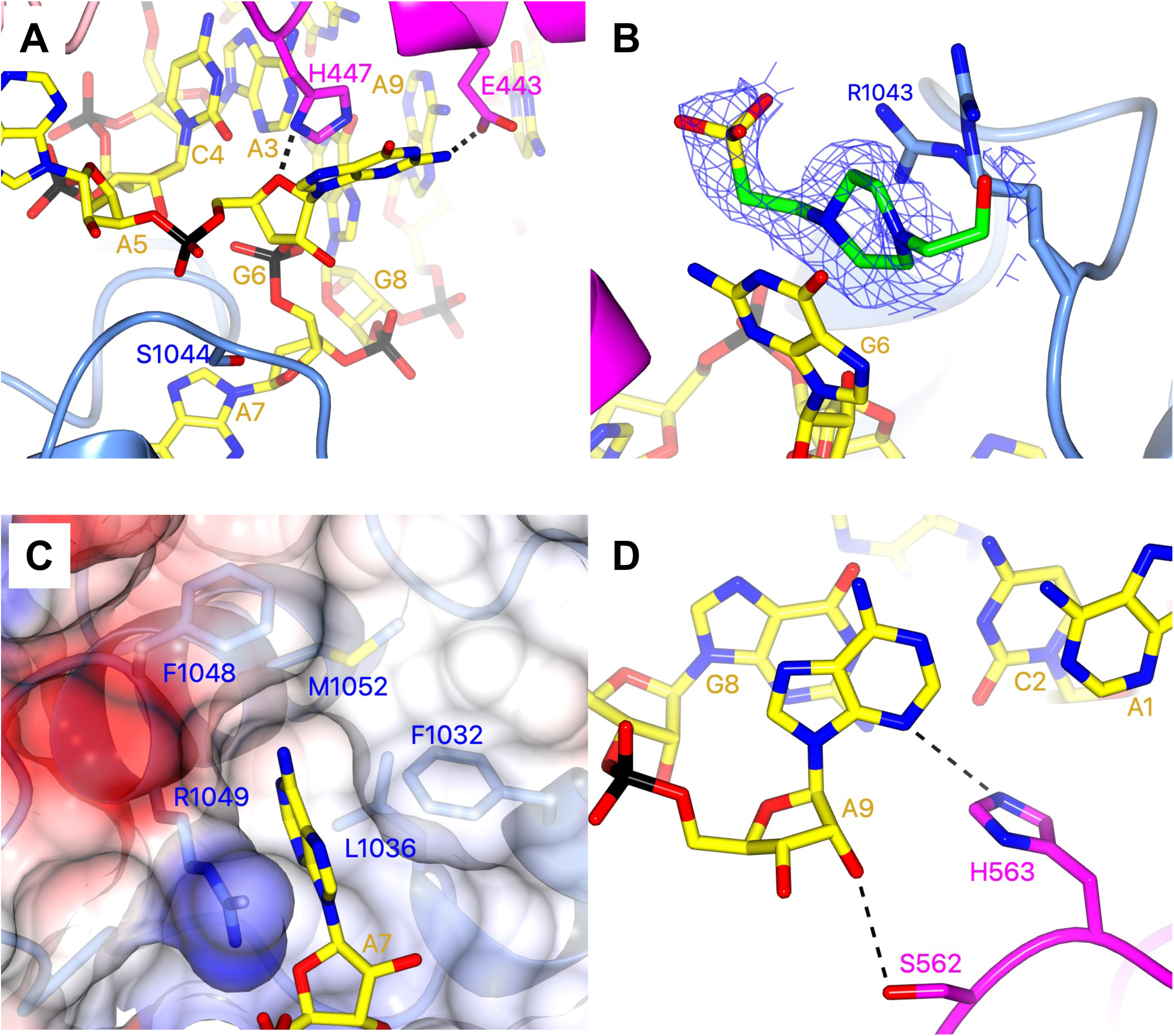
Coordination of the SFTSV 5’ RNA hook. (A) Coordination of G6 by S1044 and H447. (B) Stacking of a single HEPES molecule between the G6 base and the R1043 sidechain. The shown map is clipped to the HEPES monomer and contoured at *2σ.* (C) Coordination of the A? base which sits in a relatively tight pocket delineated by several hydrophobic residues including F1032, L1036, and F1048. Surface electrostatics for these residues is also shown and coloured according to standard practice. (D) Coordination of the A9 base which is shown interacting with S562 and H563. For clarity, the next bases in the 5’ RNA are not shown. In each panel, direct interactions are shown as dotted lines.

**Supplementary Figure 9.**
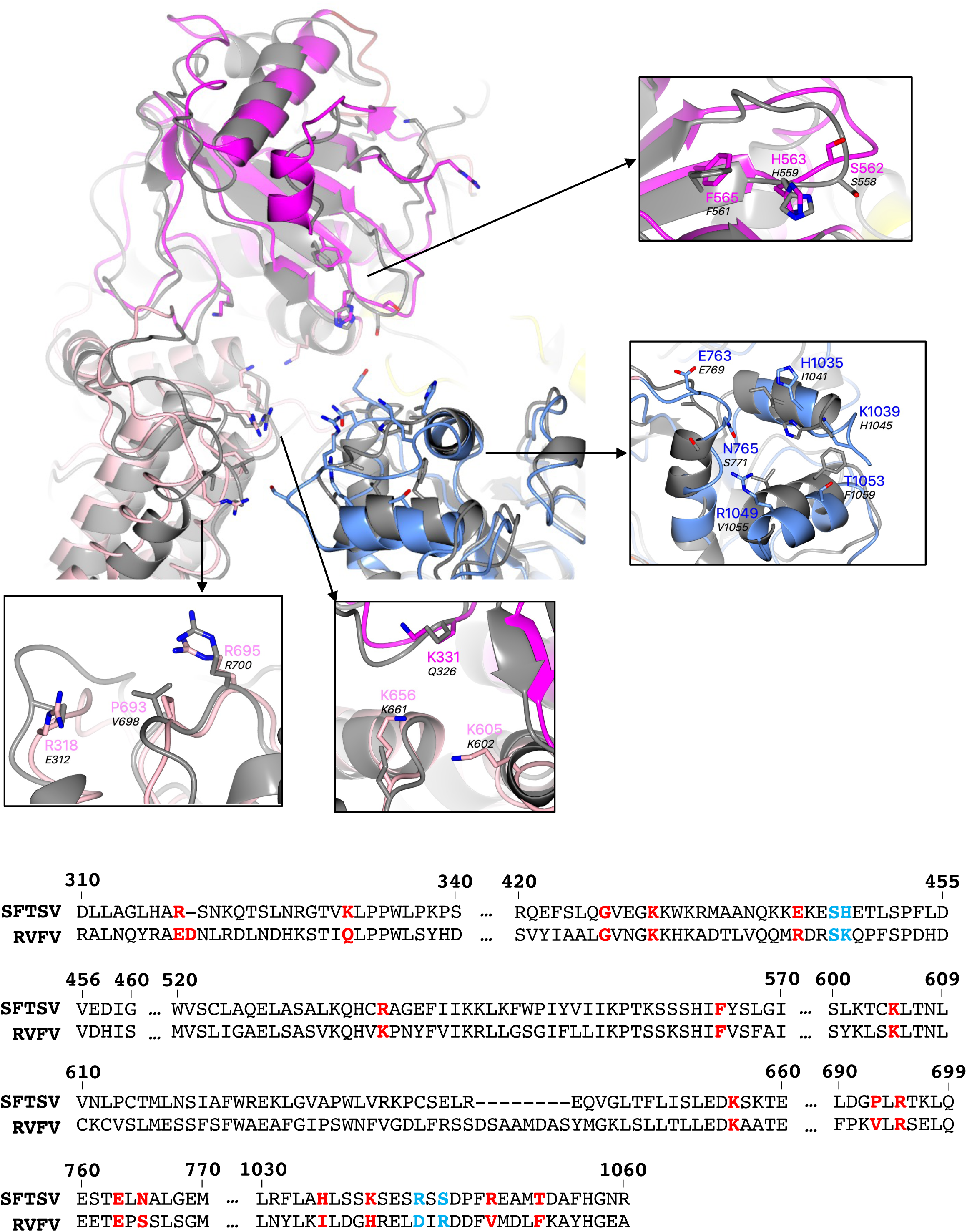
Structural superposition of the RVFV L protein onto the SFTSV L protein (hook binding site focus). The SFTSV **L** protein (5’ HOOK) is shown with the RVFV **L** protein (PDB: ?EEl, backbone coloured grey) superposed by SSM in CCP4mg [51]. Sidechains of key amino acids mutated in the cell-based mini-replicon are shown individually with close-up views. Several RVFV **L** protein amino acid sidechains appear truncated, this is because they were truncated in the ?EEl structure by the depositing authors. Residues from the SFTSV **L** protein are labelled and coloured according to the assigned domain (pink for the PA-C like-domain, blue for the fingers, and magenta for the vRBL) with the corresponding RVFV **L** protein residue labelled in black. For clarity, not all residues mutated in the cell-based mini-replicon are shown but 15 of the 23 in total are shown. A multiple-alignment diagram showing the SFTSV (UniProtKB: F1BV96) and RVFV (UniProtKB: A2SZS3) **L** protein chains generated using Clustal Omega [77] is also provided. Residues mutated in the RVFV mini-replicon system are in bold with single mutants coloured red and double-mutants coloured blue.

**Supplementary Figure 10.**
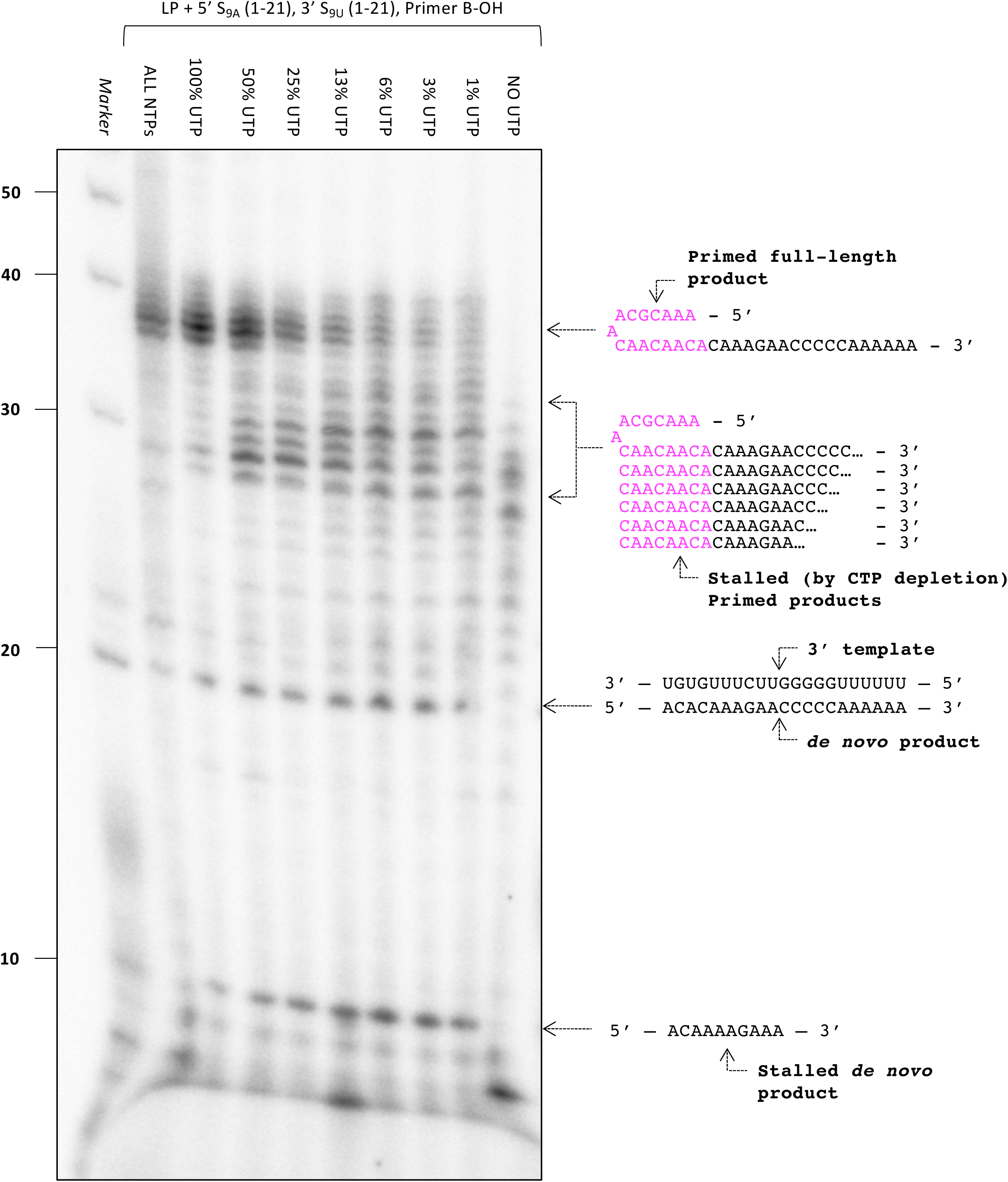
Misincorporation of UTP for CTP. The effect of removing UTP from the NTP mix in the absence of CTP on the polymerase activity of purified SFTSV L protein (LP) was tested *in vitro.* The reactions were carried out with the S_9A_ 5’ RNA (nts 1-21), S_9_u 3’ RNA (nts 1-21), and Primer B-OH present under standard polymerase assay conditions (see Materials and Methods). Products were separated by denaturing gel electrophoresis and visualised by autoradiography. Uncropped original blots/gels are provided in the Supplementary Data.

**Supplementary Figure 11.**
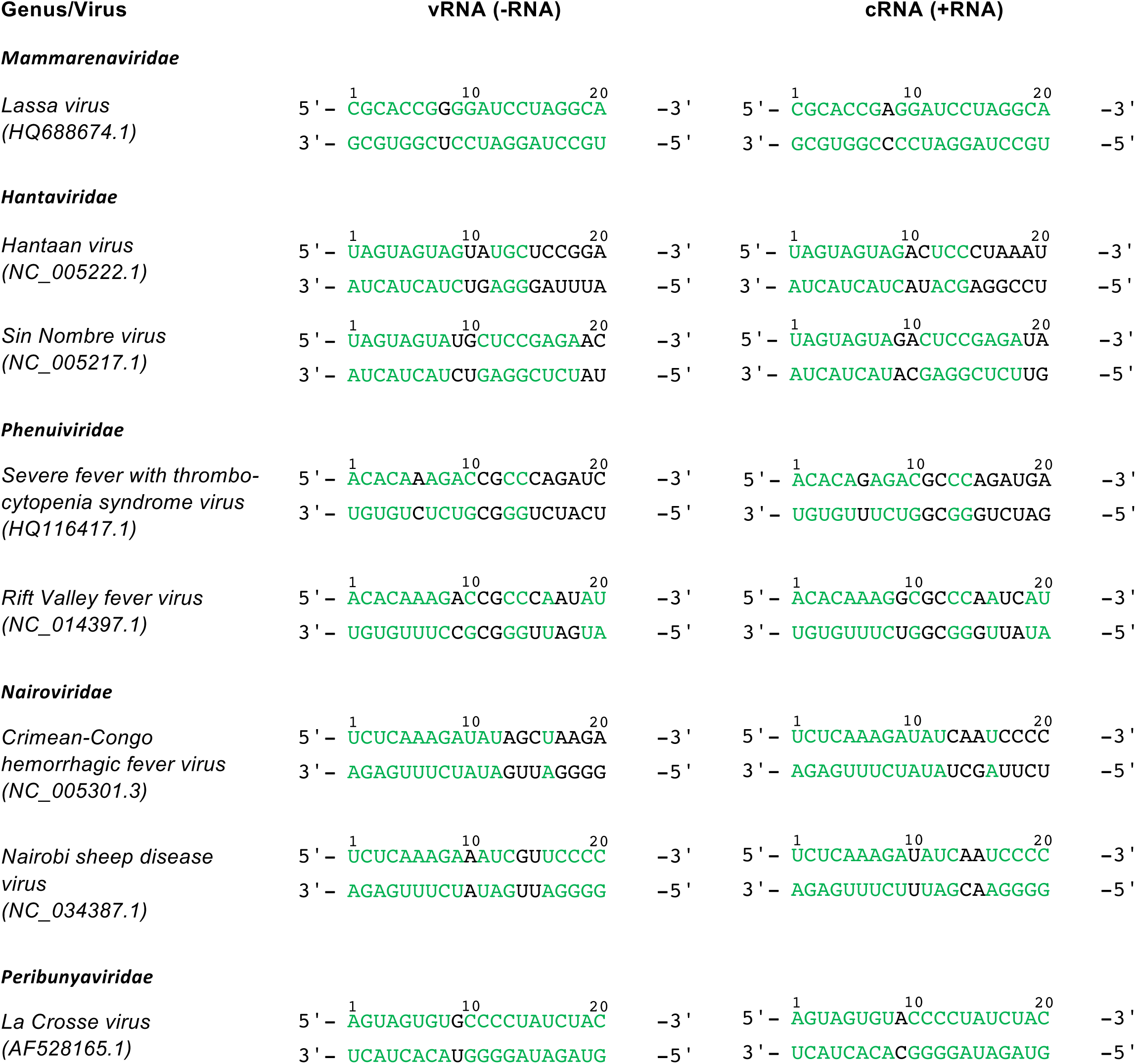
Comparison of the 5’ and 3’ L gene ends for a selection of different bunyaviruses. For each bunyavirus, the first 20 nts of the 5’ and 3’ ends of the genomic (-RNA) and antigenomic (+RNA) L gene are shown: nts coloured green are complementary, nts coloured black are not complementary. Sequences taken from the publicly-available RefSeq (NCB I Reference Sequence) Database.

**Supplementary Figure 12.**
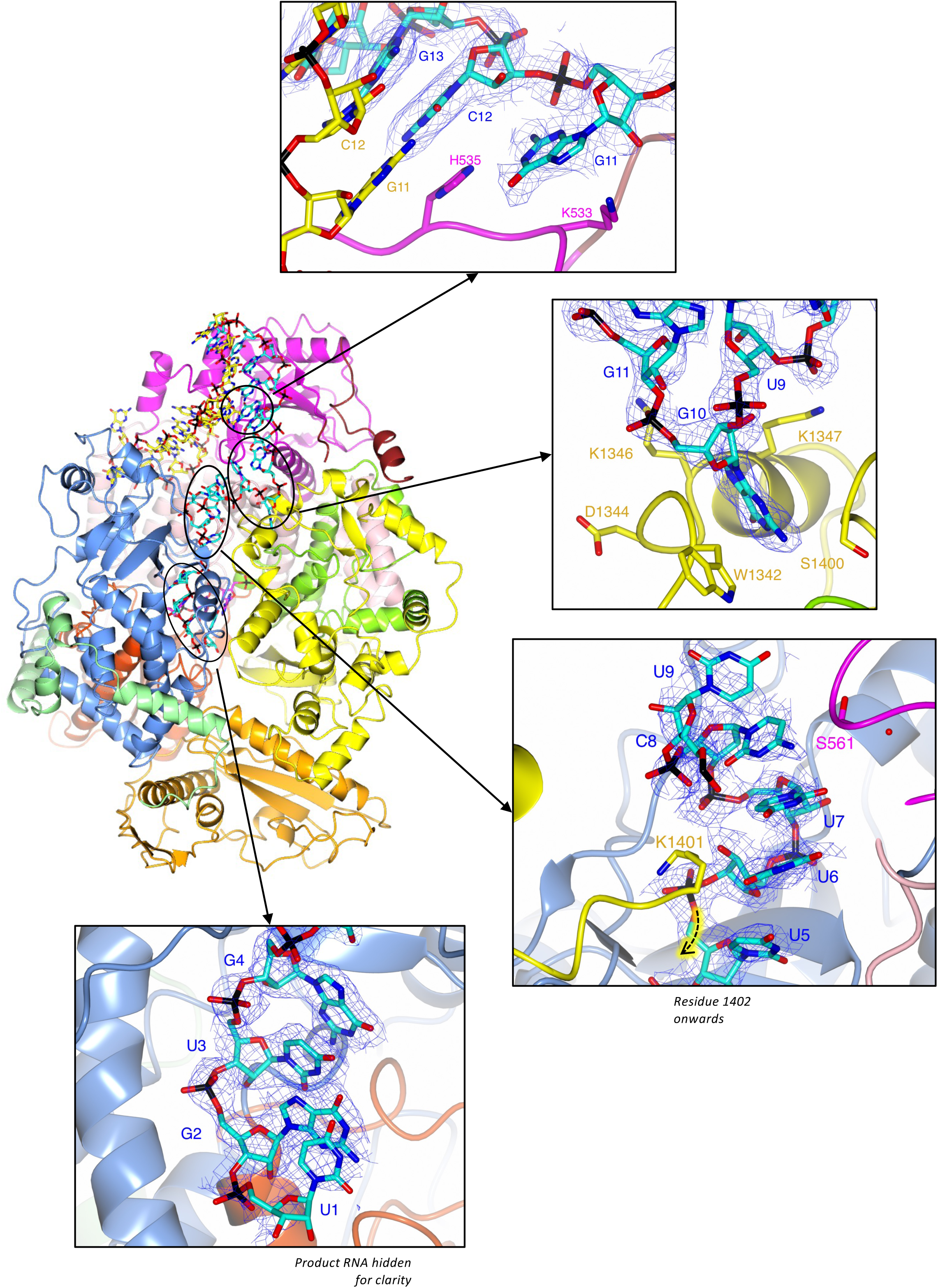
Fitting of the 3’ RNA between the distal duplex and the template entry channel. An overview of the EARLY-ELONGATION structure is shown as a ribbon with the protein in dark grey. The RNA in the EARLY-ELONGATION structure is shown as sticks and coloured yellow (5’ RNA) and cyan (3’ RNA). Zoomed-in views of the 3’ RNA in the space between the distal duplex and the template entry channel are shown in four panels. In each panel, the protein backbone and selected residues are coloured according to the protein domain: vRBL (magenta), thumb (green), thumb ring (yellow), PA-C like (light pink), and core lobe (blue). Key interacting residues and individual 3’ RNA nts are labelled accordingly. The map is shown and clipped to the 3’ RNA contoured at *4a*.

**Supplementary Figure 13.**
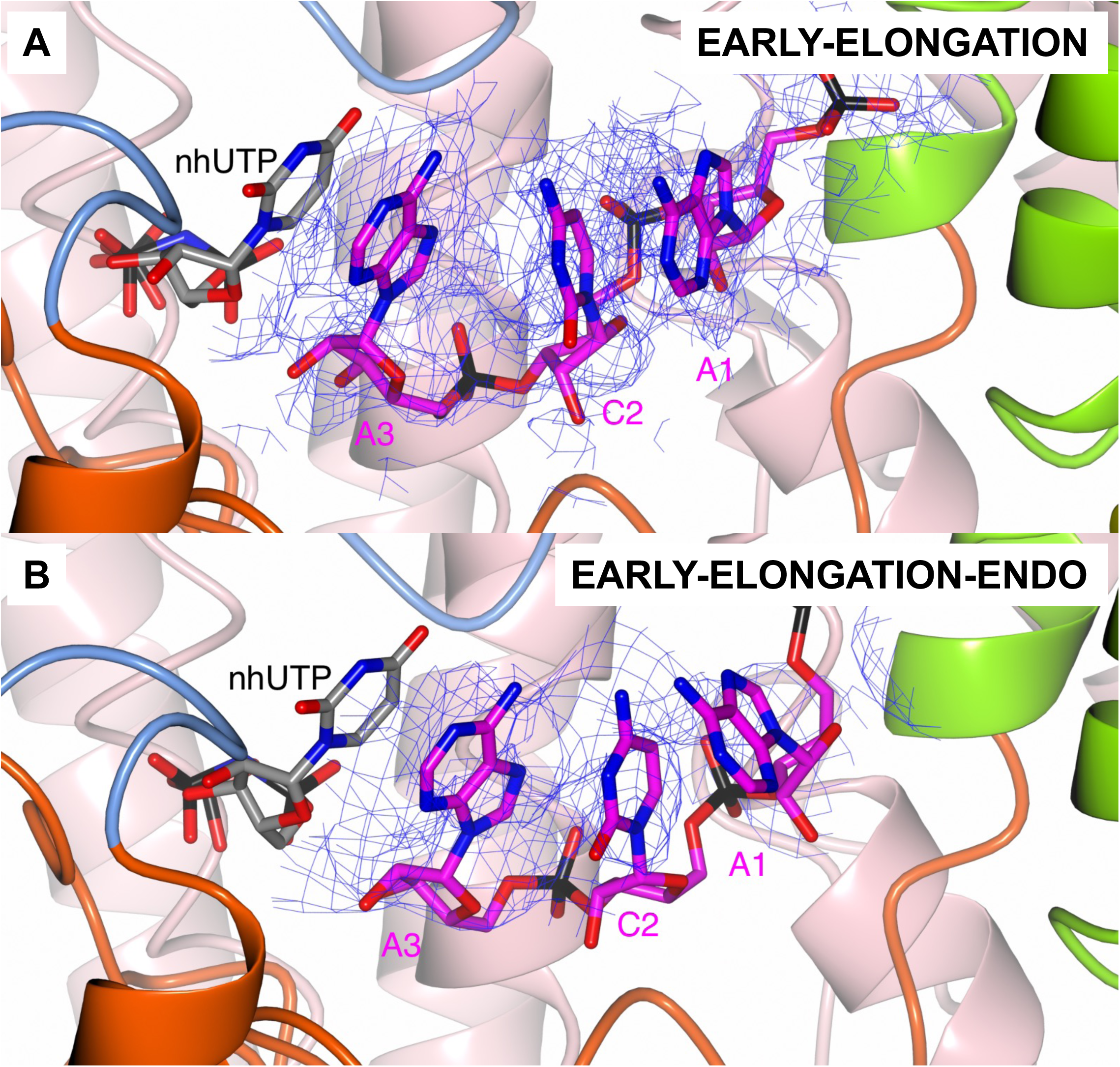
State of the product RNA at early-elongation. The product RNA is shown in the L protein core in the EARLY-ELONGATION (top) and EARLY-ELONGATION-ENDO (bottom) panels with the stalling nhUTP shown at the +1 position. For clarity, the template RNA is not shown in either panel. In both panels, the volume into which the product RNA nts have been fit is shown and is contoured at *2a*.

**Supplementary Figure 14.**
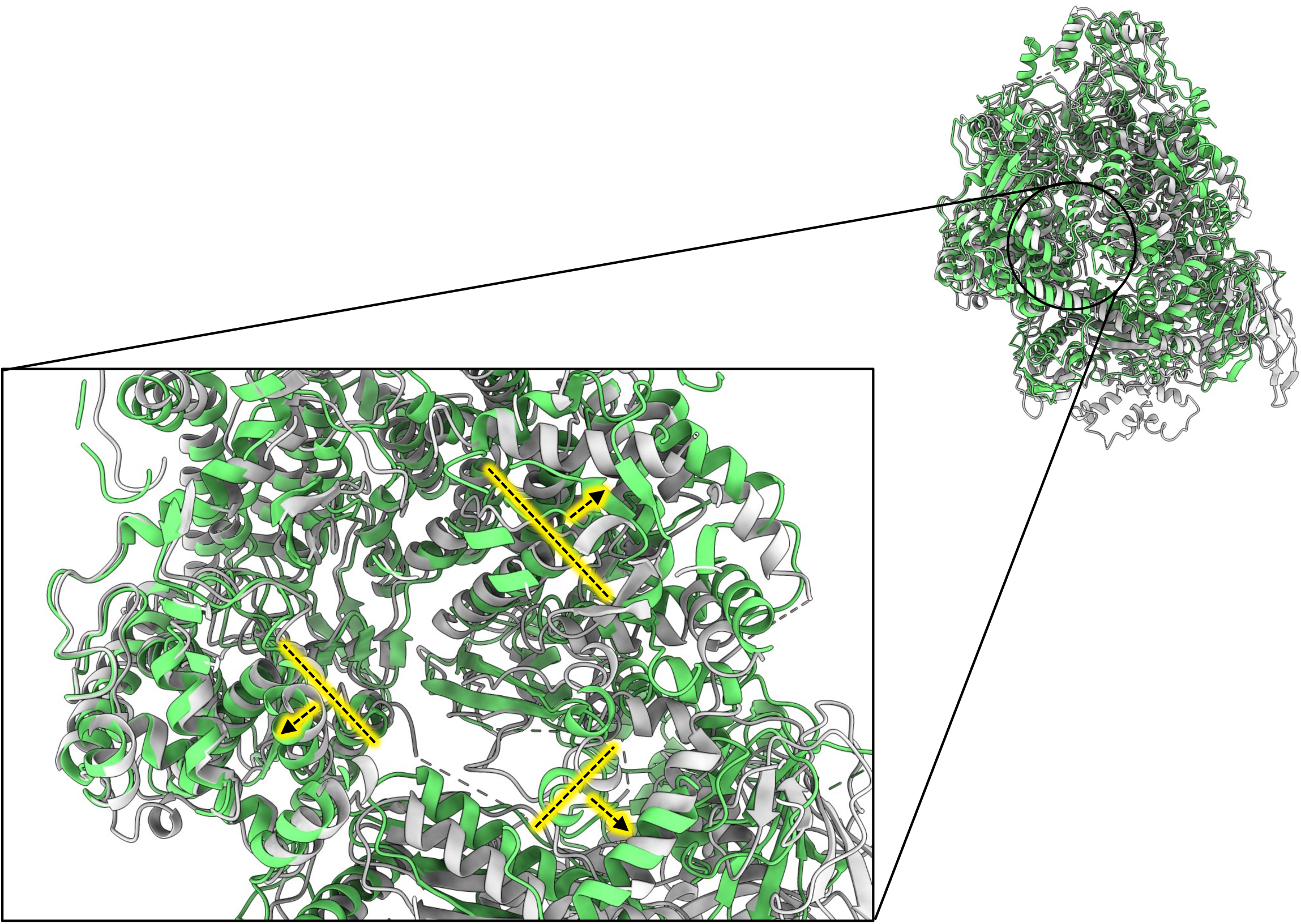
Relaxation of the L protein core from pre-initiation to late-stage elongation. The protein backbone from the published apo structure (PDB: ?ALP) and the LATE ELONGATION structures are shown in grey and green, respectively. In the zoomed-in view, dashed lines with arrows indicate the principal direction of movement from apo to late-stage elongation. For clarity, the RNA in the L protein core in the LATE ELONGATION structure is not shown.

**Supplementary Figure 15.**
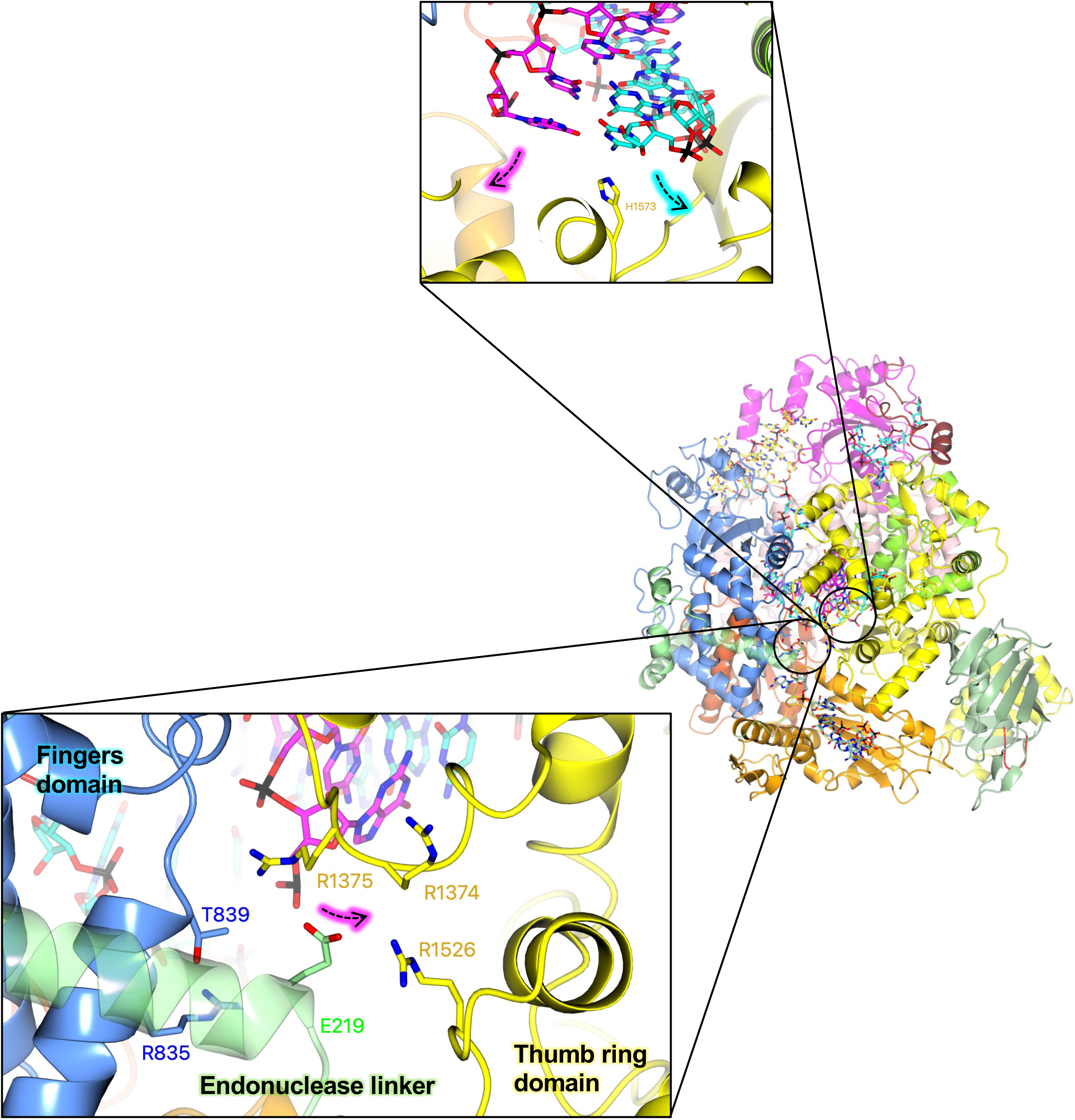
The putative SFTSV L protein product exit channel at late-stage elongation. In the top zoomed-in view, the base of the product-template duplex with the H1573 sidechain buttressing the template RNA (shown in yellow) is shown. The direction of the product (shown in magenta) are indicated by dashed arrows. In the bottom zoomed-in view, the proposed exit channel in the L protein is open under ape/resting conditions and is composed of sidechains from several L protein domains, including E219 (endonuclease-linker), T839 and R835 (PA-C like domain), R1374-R1375 and R1526 (thumb ring domain). These sidechains are shown as sticks and labelled. The path of the product RNA, which is coloured magenta, is indicated by a dashed arrow.

**Supplementary Figure 16.**
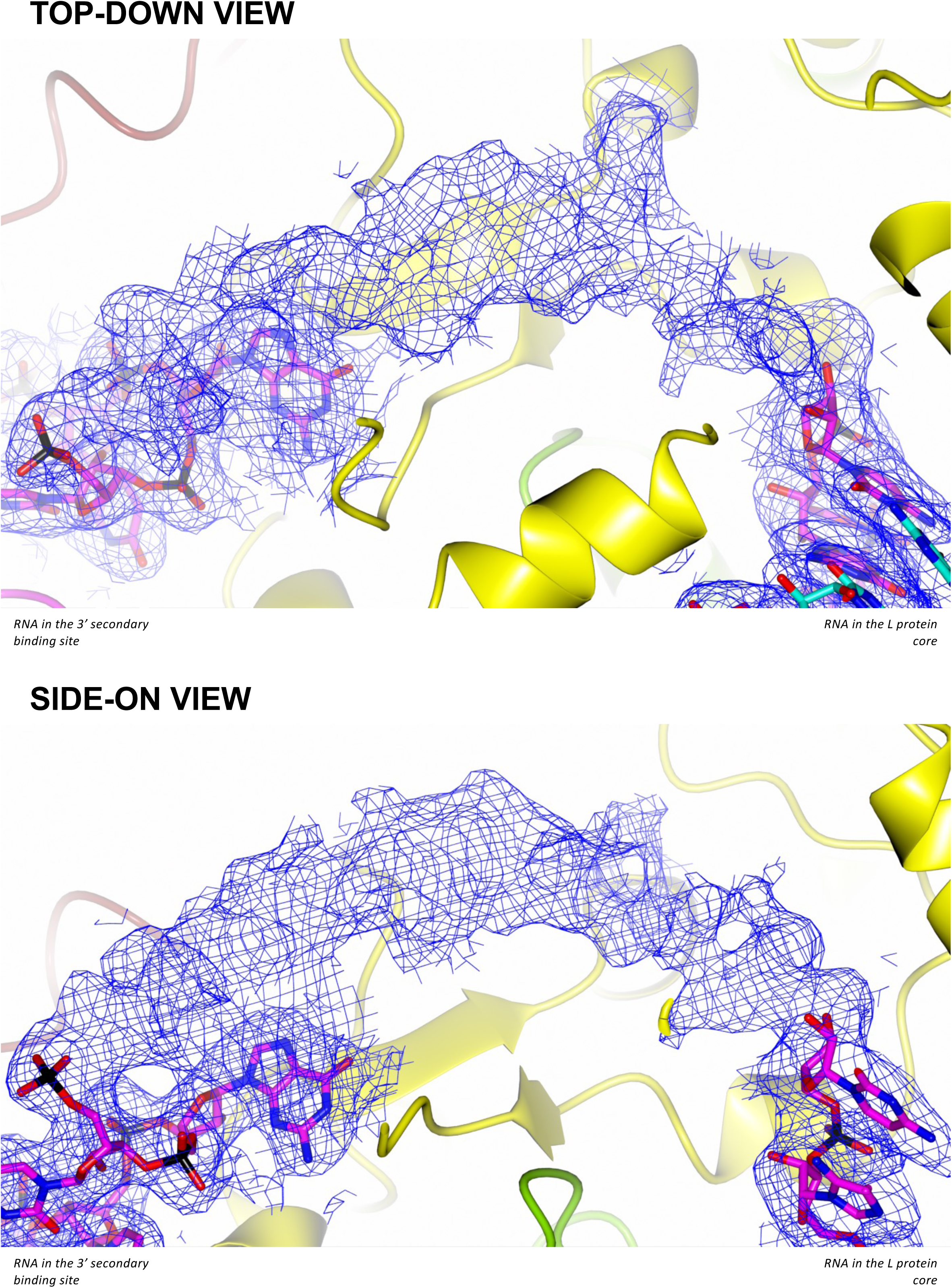
Blurred maps showing progression of the 3’ RNA towards the 3’ secondary binding site. The volume in the space between the template exit channel and the 3’ secondary binding site in the LATE ELONGATION structure is shown in a top-down (top panel) and side-on (rotated by 90°) view (bottom panel). The volume was blurred by a factor of −50 in Coot and then contoured at *4σ.* demonstrating that there is unbroken density between the 3’ RNA in the L protein core and the RNA fit in the 3’ secondary binding site.

**Supplementary Figure 17.**
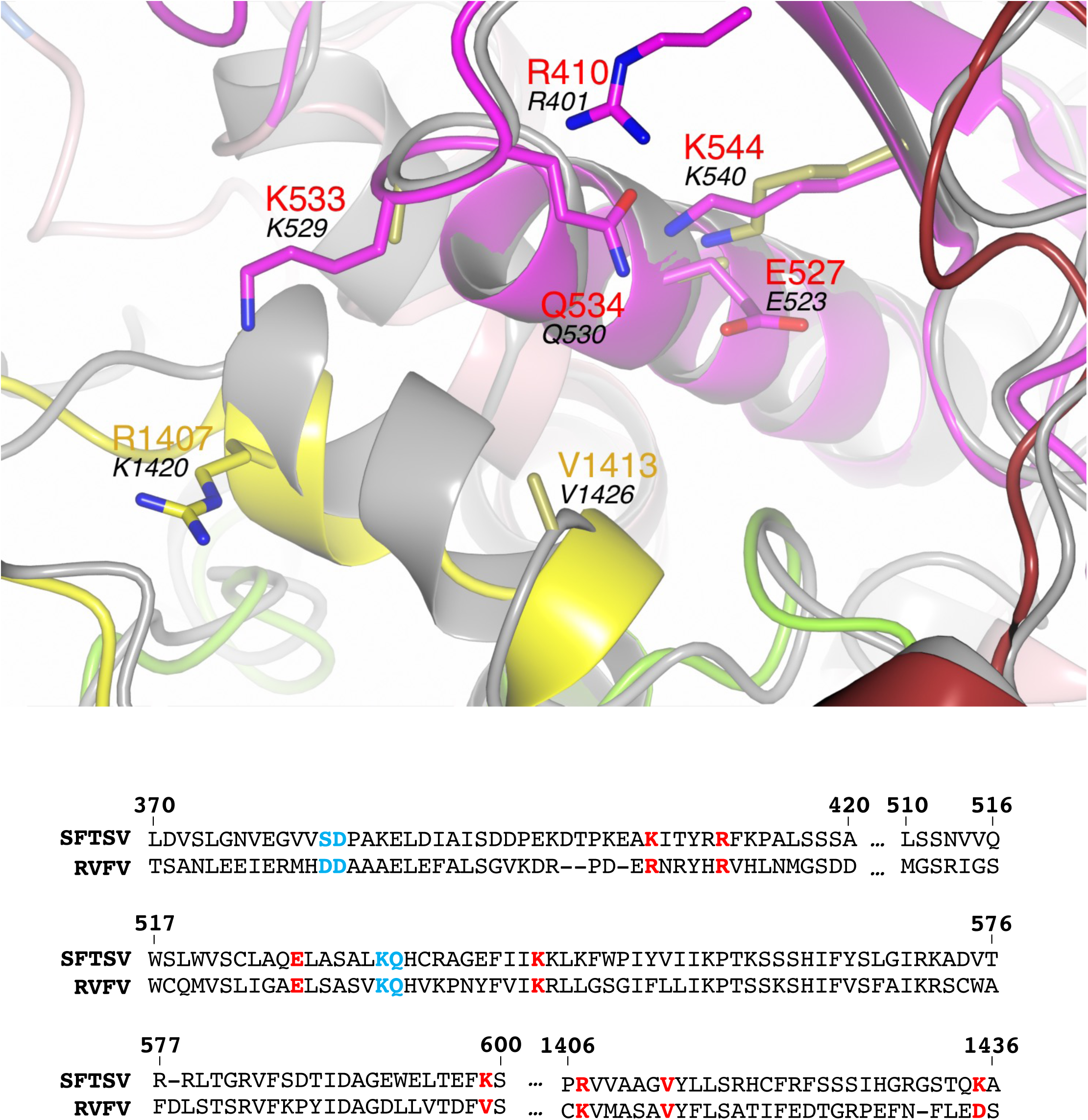
Structural superposition of the RVFV L protein onto the SFTSV L protein (focus on the 3’ secondary binding site). The SFTSV **L** protein (LATE-ELONGATION) is shown with the RVFV L protein (PDB: 7EEI, backbone coloured grey) superposed by SSM in CCP4mg [51]. Sidechains of key amino acids mutated in the cell-based mini-replicon are shown individually. Several RVFV L protein amino acid sidechains appear truncated, this is because they were truncated in the ?ALP structure by the depositing authors. Residues from the SFTSV **L** protein are labelled and coloured according to the assigned domain (pink for the PA-C like-domain, magenta/red for the vRBL, green for the thumb domain, and yellow for the thumb ring) with the corresponding RVFV **L** protein residues shown in black text. For clarity, not all the residues mutated in the cell-based mini-replicon are shown but only 7 of the 10 in total. Created using CCP4mg. A multiple-alignment diagram showing the SFTSV (UniProtKB: F1BV96) and RVFV (UniProtKB: A2SZS3) **L** protein chains generated using Clustal Omega is also provided. Residues mutated in the RVFV mini-replicon system are in bold with single mutants coloured red and double-mutants coloured blue.

**Supplementary Figure 18.**
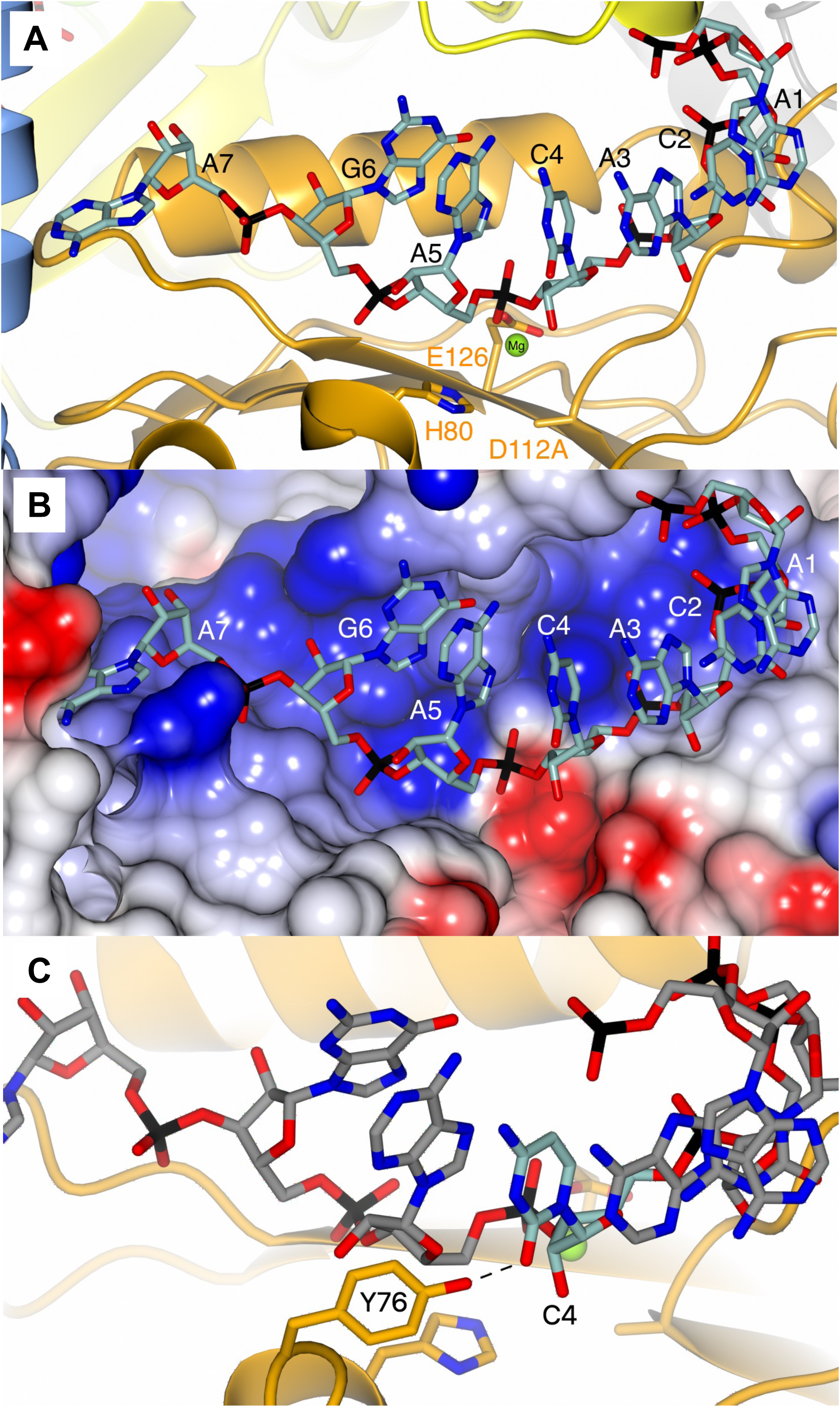
Coordination of RNA across the SFTSV L protein endonuclease domain. (A) The SFTSV L protein (LATE-ELONGATION) structure is shown focussed on the endonuclease domain. The L protein is coloured according to domain with the endonuclease, PA-C like-domain, and thumb/thumb ring domains coloured orange, blue, and yellow, respectively. The endonuclease-bound RNA is shown as cylinders and coloured sea green. Three of the main amino acid sidechains involved in catalysis are highlighted. These are H80, E126, and 0112, the latter of which has been mutated in our experiments to an alanine, hence explaining the lack of sidechain. The endonuclease-bound RNA is labelled from the 5’ end. (B) The same view as in A but with surface electrostatics overlaid showing that the RNA is bound in a positively charged cleft across the endonuclease surface. (C) A similar, zoomed, view to that shown in A demonstrating the Y76 sidechain interacting with endonuclease-bound RNA. Here, for clarity, the endonuclease-bound RNA is coloured grey with only C4 coloured sea green.

**Supplementary Figure 19.**
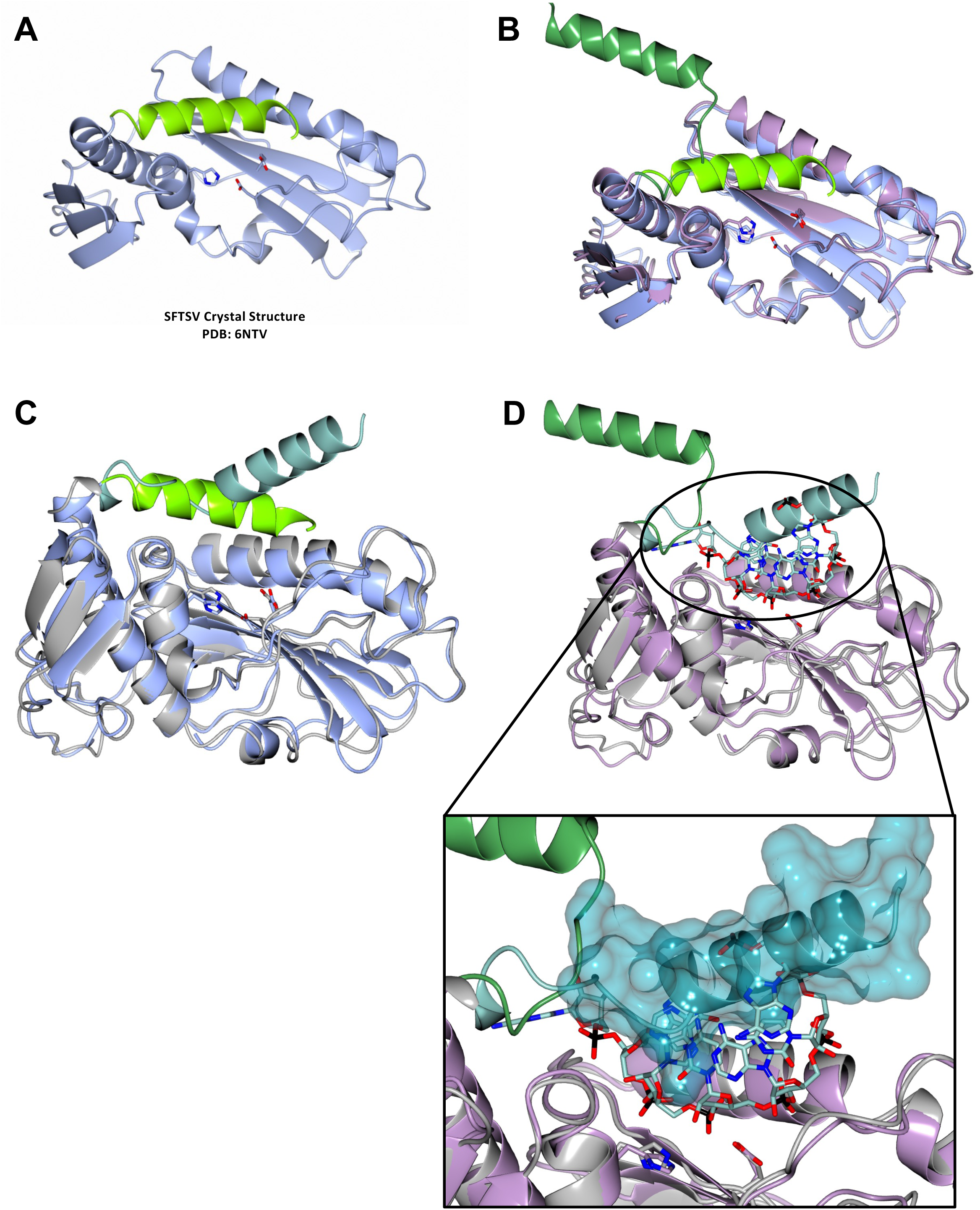
Insights into SFTSV endonuclease regulation. (A) The published SFTSV endonuclease crystal structure, PDB: 6NTV, is shown. The protein is coloured in ice blue with α helix 6 (α6, residues 211-226) shown in light green. (B) The same view as in A but with residues 1-234 of the EARLY-ELONGATION structure superposed. The EARLY-ELONGATION structure is coloured lilac with α6 (residues 211-234) shown in dark green. (C) The same view as in A but with residues 1-234 of the EARLY-ELONGATION-ENDO structure superposed. The EARLY-ELONGATION-ENDO structure is coloured dark grey with α6 (residues 211-234) shown in sea green. (D) The EARLY-ELONGATION structure is shown as in B, however, the residues 1-234 of the EARLY-ELONGATION-ENDO have also been superposed. The EARLY-ELONGATION-ENDO structure is coloured as in C. The endonuclease-bound RNA from the LATE-ELONGATION structure is also shown and is coloured yellow. A zoomed-in view of α6 is provided with the helix surface shown. In A-D, the three putative Me^2+^ coordinating sidechains (H80, D112 in the crystal structure but D112A in our cryo-EM structures, and E126) are shown and coloured according to the protein mainchain. Structures were superposed by a least-squares fit of the indicated residue ranges in CCP4mg [51].

## Notes

### Competing Interest Statement

The authors have declared no competing interest.

### Summary of Updates

This revision is to update the bioRxiv version of this paper to the latest now-accepted copy. There are only very minor changes which mostly relate to the discussion section in which we have expanded on a couple of points in response to reviewers.

